# In search of intralocus sexual conflict in the multivariate and local genetic architecture of metabolic traits in humans

**DOI:** 10.64898/2026.06.28.735027

**Authors:** Saikat Chakraborty, Kaulick Mitra, Anasuya Chakrabarty

## Abstract

Despite a common genome, males and females are remarkably different across species. Intralocus Sexual Conflict (IASC) arises when selection favours different trait values in the sexes but a shared genetic architecture constrains their evolutionary divergence. IASC is typically studied at the single-trait level, and genome-wide architectures can obscure conflict localized to specific genomic regions. We investigated IASC in the multivariate and local genetic architecture of 17 human metabolic traits and lifetime reproductive success by estimating the sex-stratified additive genetic (co)variance matrix (**G**_**mf**_) and local cross-sex-cross-trait genetic correlations. Genome-wide, between-sex covariance matrix (**B**) showed only 1.15% asymmetry, and sexually concordant (SC) genetic variation exceeded sexually antagonistic (SA) variation by 18.5-fold. Indirect evolutionary response to SC selection was 1.6-fold stronger than direct responses to SA selection. In contrast, local analyses revealed a more heterogeneous structure – among the regions with significant local correlations, 12% were consistently SA, showing opposite fitness effects in the sexes, with functional enrichment in WNT signalling, while 20% exhibited both SA and SC effects. Our results demonstrate that in human metabolic traits, SC variation predominates genome-wide, whereas SA variation is concentrated in specific local regions, highlighting the importance of integrating multivariate and local frameworks in understanding conflict.

## Introduction

Males and females share almost the entire genome, and yet they are strikingly different across taxa. The evolutionary consequences of a shared genetic architecture between the sexes represent a fundamental problem in biology and are elegantly captured by the framework of intralocus sexual conflict (IASC). IASC lies at the heart of how sexual dimorphism evolves in shared homologous traits. It arises when males and females experience sex-specific selection on shared traits and are displaced from their respective phenotypic optima [1, 2]. Under such conditions, alleles increasing fitness in one sex decrease fitness in the other due to sexually antagonistic (SA) selection acting on the same loci, creating an evolutionary tug-of-war. The intersexual or the cross-sex genetic correlation between homologous traits (r_mf_) is an estimate of the degree of shared genetic architecture between the sexes [1], and has been commonly used as a key parameter for identifying IASC [3–5]. The general idea is that a strong positive r_mf_ for a shared trait combined with opposite selection gradients on the trait in the two sexes leads to IASC. However, this framework has possible limitations – a) r_mf_ is defined for only a single shared trait between the sexes ignoring all the other correlated traits which might play important role in constraining or facilitating the evolution of dimorphism [6], and b) the relationship between r_mf_ and sexual dimorphism is not straightforward and therefore, its role in diagnosing sexual conflict is also debatable. Although early studies reported a negative correlation between r_mf_ and sexual dimorphism [7], recent theoretical as well as empirical investigations reveal that this correlation is much weaker and can also be positive [4, 5, 8]. Moreover, most of the traits of interest are highly polygenic, i.e., they are influenced by hundreds of loci. Each locus in turn affects multiple traits due to genetic correlation arising from pleiotropy or linkage disequilibrium (LD). Consequently, restricting the concept of IASC to conflict at a single locus acting on a single trait is arguably simplistic and fails to capture the full extent of multivariate genetic architecture through which sexual conflict operates [5]. A comprehensive understanding of IASC therefore requires moving beyond single-trait, single-locus perspectives towards a multivariate framework.

The intricacies of multivariate genetic architecture are statistically summarized by the additive genetic variance-covariance matrix, **G** [9]. For shared traits between the sexes, **G** takes the form of **G**_**mf**_ where it can be partitioned into four submatrices, **G**_**m**_, **G**_**f**_, **B** and **B**^**T**^ [6]:

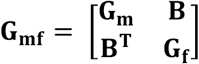

**G**_**m**_ and **G**_**f**_ are the within-sex **G** matrices and **B** is the between-sex covariance matrix. The diagonal of **B** contains the cross-sex or intersexual genetic covariances and the off-diagonals contains the cross-sex-cross-trait genetic covariances, i.e., Cov(Trait i_f,_, Trait j_m_) and Cov(Trait i_m,_, Trait j_f_). **B** is the multivariate analogue of r_mf_ and represents shared multivariate genetic architecture between the sexes [6, 10]. Often, **B** is asymmetric, i.e., Cov(Trait i_f,_, Trait j_m_) ≠ Cov(Trait i_m,_, Trait j_f_), which complicates the consequences of this shared architecture by either constraining or facilitating the evolution of sexual dimorphism depending on the direction of sex-specific selection [11–13].

Moreover, **B** represents multivariate IASC, as it not only captures the cross-sex covariances but also the cross-sex-cross-trait genetic covariances. Although termed ‘intralocus’ sexual conflict, the multivariate framework reveals that conflict operates through multiple loci affecting suites of correlated traits rather than single-locus effects on single traits. Estimating **B** to study IASC is becoming common [14–17] across several taxa and is essential in understanding the complex nature of conflict.

The canonical view of the evolution of sexual dimorphism relies on the assumption of SA selection acting on homologous traits in an initial monomorphic population, as described above [1, 18, 19]. A meta-analysis of sex-specific selection gradients while accounting for sampling error by Morrissey (2016) [20] showed that SA selection is indeed rare in nature and when present tends to be weaker compared to sexually concordant (SC) selection. More recent studies showed that evolution of dimorphism is feasible through concordant selection as much as, if not more than, SA selection [17, 21, 22]. Cheng and Houle (2020) [19] formalised this view under a multivariate framework which takes into account the direct as well as indirect effects of SA and SC selections and their effect on the evolution of dimorphism. They decomposed the **G**_**mf**_ matrix into concordant and antagonistic subspaces (**G**_**a**_ and **G**_**c**_) and the matrix **B**_**ca**_ consisiting of of differences between the male and female genetic (co)variances and the asymmetric off-diagonal components of the **B** matrix. The male-female space is mapped to the concordant-antagonistic spaces in such a way that –

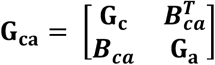

The size of the **B**_**ca**_ matrix (Frobenius norm) influence the indirect effects of SA and SC selection. They analytically showed that with high r_mf_ values, i.e., highly correlated **G**_**m**_ and **G**_**f**_, and low asymmetry in the **B** matrix, the response to SA selection is limited. Due to this high correlation between the male and female **G** matrices, the size of **G**_**a**_ relative to **B**_**ca**_ is much smaller, leading to stronger indirect response to SC selection compared to direct response to SA selection. This is particularly relevant in case of human genetic architecture where the majority of the r_mf_ values are ~ 1. This calls for a thorough investigation of the relative amount of concordant and antagonistic genetic variations and the role of indirect responses to SC selection in the evolution of sex differences in humans.

Shared genetic architecture between the sexes captured by r_mf_ and the **B** matrix represents genome-wide sexual conflict because antagonistic or concordant effects are averaged across the genome. On the other hand, dissecting the genome-wide genetic architecture into local genomic regions can help us to understand IASC with exceptional resolution. The elements of the submatrices of **G**_**mf**_ are all genome-wide estimates of genetic (co)variances and can be regarded as global genetic (co)variances/correlations. But different genomic regions can contribute disproportionately to shared genetic architecture between traits [23, 24] and sexes, and even in different directions. Importantly, even when global genetic correlations are not significantly different from zero, local genomic regions can harbour significant shared genetic variation [23]. This is especially relevant for shared traits under IASC, as the global correlations might not indicate conflict but there can be existing local conflict which we are unable to identify in a genome-wide average. Moreover, there can be local conflict resolution when global correlations represent IASC. Particularly, the local cross-sex-cross-trait genetic correlations between shared trait A in males and trait B in females can provide insight into genomic regions increasing fitness (concordant) or have opposite effects on fitness (antagonistic) in both the sexes. This would be analogous to multivariate IASC at the local level. To our knowledge, no previous study has systematically investigated how IASC is localized across the genome, examining whether genomic regions differ in their conflict status despite genome-wide concordance. Thus, understanding IASC requires integrating two complementary perspectives – a multivariate framework that captures genome-wide constraints and a local framework that identifies where conflict is concentrated across the genome.

Empirical studies of IASC originated with landmark work in *Drosophila* [3, 25] and has been extended to a wide range of taxa. In contrast, sexual conflict in humans is relatively underexplored, but the scenario is changing fast. There have been studies on IASC in humans, especially in traits such as height [26–28], though investigating conflict under a multivariate framework is rare [13, 29]. Recent advances in statistical genetics and the large-scale availability of genomic data have fundamentally advanced the scope. The development of methods that leverage both publicly available genome-wide association study (GWAS) summary statistics and individual-level data as well as whole-genome sequences has made it feasible to address quantitative genetic questions in humans in remarkable detail. Moreover, the extremely large sample sizes now available through biobanks, not only increase the precision of genetic parameter estimates but also enable detailed investigation of multivariate genetic architectures of complex traits in ways that are currently unparalleled in other species. Recent studies that have identified signatures of conflict using genomic data include – the evidence of genome-wide sex-differential selection with life history trade-off on survival and fecundity that has opposite effects on the sexes [30], signatures of SA selection [31, 32], indications of SA selection on variants affecting testosterone [33], lack of SA polymorphisms in X chromosome [34], role of recombination in resolving IASC through gene duplication [35], and identification of SA polymorphisms affecting complex diseases and traits [36]. These advances demonstrate the feasibility of rigorous IASC studies in humans using genomic data, both individual-level and publicly available summary statistics.

The choice of traits can also be decisive in shaping the asymmetry in **B** and how **B** affects sex difference, just as much as it is important in influencing the stability and structure of **G** [37]. Asymmetric **B** often indicates that one of the sexes will have unequal responses to the same selection pressure leading to the evolution of dimorphism [38]. A recent study shows that this asymmetry influences response to selection more in females than males [17]. However, it is not clear how the dynamics of asymmetric **B** works in concert with SA and SC selection and influence the evolution of dimorphism. Anthropometric traits and sex-hormones in humans of European ancestry have been shown to have a ~ 14% asymmetric **B** which acts as a constraint to the evolution of sex difference under simulated SA selection but not concordant selection [13]. Therefore, it is important to assess how **B** behaves and influences IASC for different functional categories of traits examined.

Among the many trait categories suitable for IASC investigation, metabolic traits are particularly interesting. Sex differences in metabolic and cardiometabolic traits and disorders are well documented in humans [39–41]. Energy homeostasis also shows sex-specific effects and is hypothesized to have evolved under sex-specific selection with starvation resistance and propensity to obesity evolving simultaneously in females [42]. To the best of our knowledge, the studies on sex difference in metabolic traits in humans are exclusively univariate. However, metabolism is composed of networks of traits involved in energy storage and allocation, glucose and insulin regulation, hepatic metabolism, cardiovascular regulation, inflammation, etc. Therefore, for in-depth studies of sex differences in metabolic traits, we need a multivariate approach. Moreover, traits closely linked to metabolism such as BMI, waist-to-hip ratio, etc., show sex-specific fitness associations [13, 28] which are indicative of potential SA genetic variation, making metabolic traits as ideal candidates for investigating multivariate IASC in humans.

In this study, we investigated intralocus sexual conflict (IASC) across 17 metabolic and physiological traits along with lifetime reproductive success (LRS) as a fitness proxy in contemporary humans at different levels using publicly available GWAS summary statistics –

a. We estimated the multivariate genetic architecture of these 18 traits by estimating the full **G**_**mf**_ and ascertained the asymmetry in **B**. The correlations in **G**_**mf**_ are regarded as global genetic correlations.
b. We mapped the male-female space (**G**_**mf**_ matrix) to concordant and antagonistic spaces to estimate the size of concordant and antagonistic genetic (co)variation, and estimated the direct and indirect respondabilities to SC and SA selection in response to random selection gradients.
c. We examined whether the **B** matrix constrains or facilitates the evolution of sexual dimorphism in these traits under random SC and SA selection.
d. To determine whether local antagonistic architecture persists despite genome-wide concordance, we estimated the local cross-sex and cross-sex-cross-trait genetic correlations of the traits which are significantly correlated to fitness (LRS) in each sex and investigated local IASC.
e. Finally, we evaluated the regions which are predominantly concordant or antagonistic, annotated them to map the associated genes, and identified enriched biological pathways in those regions.

## Methods

### a) Datasets and GWAS summary statistics

We used publicly available sex-stratified GWAS summary statistics for all the 17 metabolic and physiological traits and LRS from the Neale lab webpage (https://www.nealelab.is/uk-biobank). The GWAS were performed on individuals of white-British ancestry in the UK Biobank. The male and female sample sizes for the traits ranged from 147,044 to 193,953. The exact sample sizes are provided in the **Supplementary Table 1**. LRS was used as a fitness proxy and is measured by the number of children sired in males and number of live births in females. The remaining 17 traits comprise of a panel of metabolic and physiological biomarkers, including markers of glucose and lipid metabolism – glycated haemoglobin (HbA1c), low-density lipoprotein cholesterol (LDL), high-density lipoprotein cholesterol (HDL), and triglycerides (TRI); renal function biomarkers – creatinine (CRE) and urea (URE); hepatic biomarkers – alanine transaminase (ALT), aspartate aminotransferase (AST), alkaline phosphatase (ALP), gamma-glutamyl transferase (GGT), albumin (ALB), bilirubin (BIL); cardiovascular regulation – systolic blood pressure (SYS), diastolic blood pressure (DIA), mineral homeostasis – calcium (CAL); and endocrine growth and metabolic signalling – insulin-like growth factor 1 (IGF-1).

**Table 1.** Classification of concordant and antagonistic local genomic regions based on the score *S* which considers the sign of the local cross-sex-cross-trait genetic covariances, and the signs of the genome-wide sex-specific genetic covariance between the traits and LRS. There are 8 different combinations of the signs, and the classifications and interpretations are given accordingly.

|  | **Cov (Trait $A_m$, Trait $B_f$)** | **Cov (Trait $A_m$, LRS$_m$)** | **Cov (Trait $B_f$, LRS$_f$)** | **$S$** | **Classification** | **Interpretation** |
| --- | --- | --- | --- | --- | --- | --- |
| 1) | + | + | + | + | Concordant | Shared alleles increase fitness in both the sexes |
| 2) | + | - | - | + | Concordant | Shared alleles decrease fitness in both the sexes |
| 3) | - | + | + | - | Antagonistic | Shared alleles have opposite effect on fitness in both the sexes |
| 4) | - | - | - | - | Antagonistic | Shared alleles have opposite effect on fitness in both the sexes |
| 5) | + | - | + | - | Antagonistic | Shared alleles have opposite effect on fitness in both the sexes |
| 6) | - | - | + | + | Concordant | Shared alleles increase fitness in both the sexes |
| 7) | + | - | + | - | Antagonistic | Shared alleles have opposite effect on fitness in both the sexes |
| 8) | - | + | - | + | Concordant | Shared alleles increase fitness in both the sexes |

All the traits except LRS were normalized at the source before performing GWAS using inverse-rank normal transformation. GWAS was performed separately for males and females for each trait using age, age^2^, and principal components 1-20 to control for population stratification (https://www.nealelab.is/uk-biobank). The GWAS summary statistics data primarily used in analyses consisted of the SNP ID (rsID), chromosome, position (in base pair), effect size (slope of the GWAS – beta), standard error of effect size, P-value of the effect size, effect allele, other allele, minor allele frequency (MAF), and INFO (a parameter indicating the SNP imputation quality).

### b) Quality control of GWAS summary statistics

We restricted our analyses to HapMap3 [43] SNPs in this study, which have a high imputation quality. We performed standard QC of the GWAS summary statistics by removing strand ambiguous and duplicate SNPs, filtering SNPs for a MAF > 0.01 and INFO > 0.9. 1000 Genomes Phase 3 European (EUR) [44] LD reference panel was used for estimating both global and local genetic correlations. We removed the extended major histocompatibility (MHC) region on chr 6: 25 Mb - 35 Mb from all the GWAS summary statistics due to the presence of long-range LD. Such complicated LD patterns might affect the correlation estimates as the methods used are largely dependent on LD between SNPs.

### c) Estimating global genetic correlations in G_**mf**_

We estimated all the four submatrices of **G**_**mf**_, the within-sex cross-trait genetic (co)variances and correlations – **G**_**m**_ and **G**_**f**_, and the cross-sex and cross-sex-cross-trait genetic covariances and correlations – **B** and **B**^**T**^ for all the 18 traits in males and females. All these covariances, correlations, and SNP-heritabilities were genome-wide estimates (global) estimated using cross-trait Linkage Disequilibrium Score-Regression method (LDSC) [45, 46]. Under this model, genetic covariance between two traits is the slope of the regression of the product of Z-scores of the same SNPs from two studies on LD-scores [45] (**Supplementary Note**). The genetic correlation between *trait 1* and *trait 2* (or males and females) was calculated as 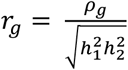, where ρ_g_ is the genetic covariance between traits 1 and 2 (males and females), and h_1_^2^ and h_22_ are the heritabilities of the two traits (or two sexes).

We applied Benjamini-Hochberg false discovery rate (FDR < 0.05) correction for multiple testing to get adjusted P-values across 630 unique correlations included in the **G**_**m**_, **G**_**f**_, **B**, and **B**^**T**^.

### d) Estimating concordant and antagonistic genetic variation and respondabilities

An important aspect of male-female genetic (co)variation is the ability to respond to concordant and antagonistic selections. The amount of concordant and antagonistic genetic variation and the covariation between them can potentially influence whether sex differences will evolve and at what rate. To disentangle these two kinds of variation in human metabolic traits, we transformed the male-female trait space to concordant-antagonistic trait space following the framework proposed by Cheng and Houle (2020) [19]. The transformed variables are the functions of trait means of the two sexes and the mean difference between the sexes.

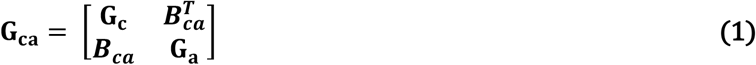

The matrix **G**_**c**_ summarises the genetic variation in the concordant subspace which allows for evolutionary response at the same rate and direction to selection in both the sexes. The genetic variation in the antagonistic subspace is captured by **G**_**a**_ which allows for evolutionary response in opposite directions to selection. **B**_**ca**_ is the matrix of genetic covariation that affects the indirect responses to selection, e.g., indirect antagonistic response to concordant selection. The relationship between the **G**_**ca**_ matrix and the male-female space can be given by the equation –

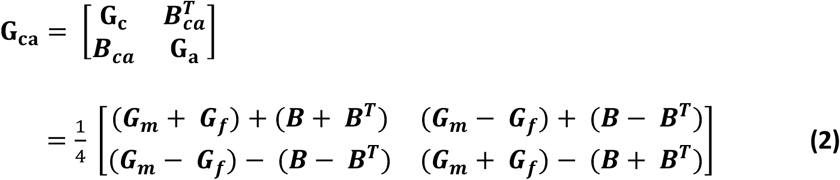

The size of the matrices **G**_**a**_ and **B**_**ca**_ given by their Frobenius norms (∥ ***G***_***a***_ ∥ and (∥ ***B***_***ca***_ ∥) influence the average rate of change of sexual dimorphism under antagonistic and concordant selection. Concordant selection indirectly contributes through **B**_**ca**_ (covariation between concordant and antagonistic variation) to the evolution of sex differences. Cheng and Houle showed that the size of **B**_**ca**_ depends on the relative size of **G**_**a**_ and **G**_**c**_, e.g., if ∥ ***G***_***a***_ ∥≪ ∥ ***G***_***c***_ ∥, then ∥ ***B***_***ca***_ ∥≥ ∥ ***G***_***a***_ ∥. This condition is particularly relevant when ***G***_***m***_ ≈ ***G***_***f***_, which is probable in humans given the high r_mf_s. If ∥ ***B***_***ca***_ ∥≥ ∥ ***G***_***a***_ ∥, then indirect responses to concordant selection will be large enough to control the evolution of dimorphism rather than direct responses to antagonistic selection.

Another metric to evaluate how much of the response to concordant and antagonistic selection falls within the concordant and antagonistic subspace is respondability [19, 47] to random skewers [48, 49]. It is the total length of the vector of responses to a selection vector of unit length.

Respondabilities to concordant and antagonistic selection vectors is given by –

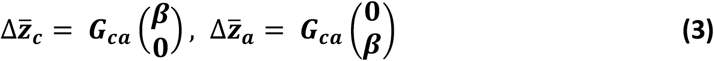

We algebraically estimated the **G**_**ca**_ using equation (2), and then calculated the Frobenius norms of the sub-matrices to evaluate the amount of concordant and antagonistic genetic (co)variation. We used the random skewers method as described by Cheverud and Marroig (2007) [49] to calculate the direct and indirect respondabilities to 1000 random concordant and antagonistic selection vectors using equation (3) [19].

We also calculated a metric for matrix similarity (S) which is a scalar measure of matrix correlation following Cheng and Houle (2020). For matrices **F** and **M** –

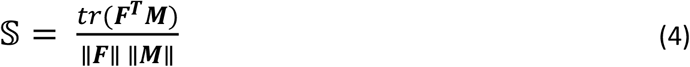

Where tr() is the trace and ∥•∥ is the Frobenius norm. S is constrained to fall between 1 and −1, where 1 denotes perfect positive correlation and −1 denotes perfect negative correlation, while 0 denotes no relationship. We calculated S between **G**_**m**_ and **G**_**f**,_ **G**_**m**_ and **B, G**_**f**_ and **B**, and **B** and **B**^**T**^.

We adapted the SAS code available from Cheng and Houle (2020) [19] and performed the entire analyses in R [50] (4.6.0).

To include sampling errors of estimates, we used 200 leave-one-out block jackknife pseudovalues for genetic (co)variances from LDSC. The jackknife values were estimated by dividing the genome into 200 blocks and then estimating (co)variances leaving each block out at a time. We built a jackknife distribution of 200 **G**_**mf**_ matrices, and performed the above analyses on each one of them. We reported the point estimates from the original matrices and 2.5% and 97.5% jackknife quantiles for the matrix norms and similarities. For respondabilities, we reported the median, 2.5% and 97.5% quantiles from the distribution of 1000 response vectors generated using random skewers.

### e) Investigating asymmetry in B and the role of B in constraining the evolution of sexual dimorphism in these traits

#### i) Asymmetry in B

Asymmetry in **B** is influenced by sex-specific selection and indicates sex difference in the multivariate genetic architecture of traits [11]. To ascertain the degree of asymmetry in **B**, we decomposed **B** into its symmetric and skew symmetric components[11]. An asymmetric matrix **A** can be decomposed into the symmetric and skew-symmetric components[51], and the matrix **A** can be represented as a sum of these two components:

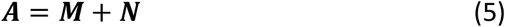

where **M** = ½(**A**+**A**^T^) and **N** = ½(**A**-**A**^T^) are the symmetric and skew-symmetric components respectively, such that **M** = **M**^T^ and **N** = −**N**^T^. We can also show that the total sum of squares can be partitioned into independent symmetric and skew-symmetric components [51], and the proportion of the matrix **A** that is symmetric or skew-symmetric can be determined. We estimated the proportion of **B** that was asymmetric from the skew-symmetric component without the diagonal. We used the *asymmetry* package[52] in R for this matrix decomposition.

#### ii) B as a genetic constraint or facilitator under concordant or antagonistic selection

**B** portrays the shared genetic architecture between the sexes and can both constrain or facilitate the evolution of sexual dimorphism. This depends on the extent of asymmetry in **B** and the direction of sex-specific selection – SC or SA. To investigate whether **B** acts as a constraint predominantly under SA selection compared to SC, we used the multivariate breeder’s equation for shared traits between sexes [6] in combination with the random skewers framework adapted from Cheverud (1996) [48] and Cheverud and Marroig (2007) [49]. We simulated random SC and SA selection vectors and quantified the evolutionary responses. Because the simulated selection vectors span a broad range of possible selection directions, some may not correspond to biologically relevant selection regimes. However, our objective was to investigate the evolutionary consequences of the observed multivariate genetic architecture of human metabolic traits rather than estimating contemporary selection. Random skewers analyses are well suited to this purpose because they evaluate the response of the covariance structure across a wide range of potential selection directions.

For two sexes, the breeder’s equation can be represented as[6]:

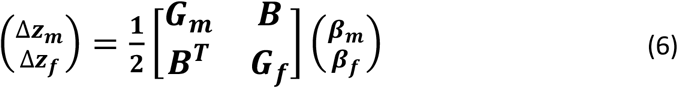

Where **Δz**_**m**_ and **Δz**_**f**_ are the short-term predicted evolutionary responses in males and females respectively, **G**_**mf**_ is the multivariate genetic architecture in both the sexes, and **β**_**m**_ and **β**_**f**_ are the vector of linear selection gradients in males and females. The equation is multiplied by ½ to adjust for the equal genetic contribution of both the parents to the offspring. To estimate selection responses without the contribution of the shared genetic architecture, **B** can be constrained to zero.

In such a scenario, we can write equation (3) as

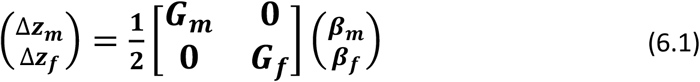

Now, if we multiply **G**_**mf**_ with the same random skewers **β**_**m**_ and **β**_**f**_, and compare the evolutionary responses **Δz**_**m**_ and **Δz**_**f**_, any difference in the responses will be due to the differences in the **G**_**mf**_.

Based on this principle, selection skewers were sampled from a multivariate normal distribution with mean zero and covariance matrix equal to the identity matrix and then normalized to unit length. The same selection vectors were used for males and females (**β**_**m**_ and **β**_**f**_), 1000 for each sex, to simulate SC selection. We also simulated SA selection by reversing the direction of the random selection vectors for males and females (<**β**_***m***_**β**_***f***_ = 180◦). This was an extreme scenario where the selection on males and females is exactly in the opposite direction.

We first projected **β**_**m**_ and **β**_**f**_ onto **G**_**mf**_ without **B** (equation 6.1) and calculated the angle between male and female responses, **Δz**_**m**_ and **Δz**_**f**_ as a measure of vector correlation. Then we included **B** (equation 6) and again measured the angle between the responses. Larger angles (θ) between **Δz**_**m**_ and **Δz**_**f**_ indicate greater evolutionary divergence between sexes. If **B** acts as a constraint, it will decrease the angle, θ, between male and female responses by making males and females more similar to each other. [13, 21, 53].

The angle θ was defined as –

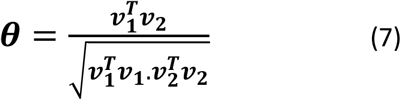

where **v**_**1**_ and **v**_**2**_ are the two vectors and **θ** is the angle between them.

### f) Genetic correlation with LRS

Within the **G**_**mf**_ framework, we estimated genetic correlations between the 17 metabolic and physiological traits and LRS in males and females, resulting in 34 sex-specific trait-fitness correlations. Traits exhibiting statistically significant genome-wide genetic correlations with LRS after multiple-testing correction in at least one sex were retained for downstream analyses. Local cross-sex and cross-sex-cross-trait genetic correlations were subsequently examined only for these traits, thereby focusing on genomic regions underlying traits with evidence of fitness relevance as IASC requires shared traits to influence fitness.

### g) Estimating local genetic correlations

We estimated local cross-sex and cross-sex-cross-trait genetic correlations for the traits in males and females which showed significant trait-fitness correlations in the step above. We used the ρ-HESS framework to estimate these local genetic correlations [23], which decomposes genome-wide genetic covariance into approximately independent LD-defined genomic regions, using GWAS summary statistics. Within each region, local SNP-heritability, genetic covariance, and genetic correlation were estimated for each trait combination. As the cross-sex and cross-sex-cross-trait genetic correlations influence IASC, and local correlations are computationally intensive, we did not extend the framework to local cross-trait genetic correlations within sexes. A total of 1691 LD-blocks, each of length 16.4 Mb were analysed, and multiple testing correction was applied using a Bonferroni adjusted threshold of P < 2.96 × 10^−5^ (0.05/1,691). Loci with P-value below this threshold were considered to exhibit significant local genetic correlation and were subsequently visualized using Manhattan-styled plots.

### h) Classification into antagonistic and concordant regions

We defined sexual antagonism/concordance at the local, multivariate level by integrating the signs of significant local cross-sex-cross-trait genetic covariance with sex-specific fitness associations using the following rule –

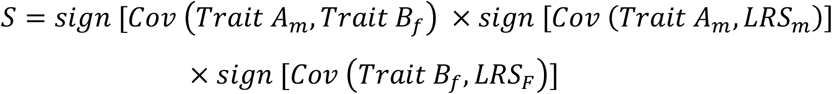

Regions with *S* = +*ve* were classified as concordant where genetic variation has aligned fitness effects across sexes, whereas regions with *S* = −*ve* were classified as antagonistic, which harbour alleles with opposing fitness consequences in males and females. This rule combines the direction of the local cross-sex-cross-trait genetic covariance with the sex-specific genome-wide trait-fitness genetic covariances. This framework extends single-trait IASC detection to multivariate genetic architecture at local genomic scales. For example, when traits A and B are similarly associated with LRS in their respective sexes, a negative cross-sex-cross-trait genetic covariance between them indicates an antagonistic region, as the region increases trait A in one sex and decreases trait B in the other, but decreasing trait B means decreasing LRS in that sex. Hence, the local genomic region has opposite effects on fitness in the sexes mediated through a local cross-sex-cross-trait genetic covariance. The detailed classification based on the score S is given in Table 1.

Based on our classification regime, we identified and flagged regions which were predominantly concordant, predominantly antagonistic, and mixed (half of the time concordant and half of the time antagonistic) category. We proceeded with annotating one region from each of the above three categories. As there were multiple trait combinations, we chose the trait with the highest local SNP-heritability in the specific region and proceeded with downstream analyses.

### i) Gene mapping and pathway analyses

We annotated the SNPs for the trait with highest SNP-heritability to map the associated genes in those regions. We used positional (± 10 kb windows), eQTL (GTEx v 8), and chromatin mapping as implemented in FUMA (v1.8.2) [54] to map the genes. We used the mapped genes from the three categories to perform pathway analyses to identify biologically enriched pathways in predominantly concordant, predominantly antagonistic, and mixed regions using Enrichr [55, 56]. Enrichr is a meta-database which includes curated pathways from several other databases.

## Results

### a) Genome-wide genetic correlations (G_mf_)

#### i) Heritabilities, cross-trait and cross-sex genetic correlations

We estimated the full **G**_**mf**_ covariance and correlation matrices along with the four submatrices using LDSC regression (**Figure 1**). All the traits showed significant heritabilities in both the sexes with sex-specific heritabilities in LRS, systolic (SYS) and diastolic (DIA) blood pressures (**Supplementary Figure 1**). Out of a total of 630 genetic correlations in **G**_**mf**_, 349 were significant at FDR < 0.05. The within-sex male and female cross-trait correlations were moderate to high (**Supplementary Tables 2 & 3**). The highest male genetic correlation was between the liver enzymes ALT and AST (0.70 ± 0.02) and the lowest correlation was between TRI and IGF1 (−0.08 ± 0.03). On the other hand, the highest female correlation was between systolic (SYS) and diastolic (DIA) blood pressures (0.68 ± 0.02), while the lowest one was between SYS and bilirubin (BIL) (−0.07 ± 0.03). Among the male cross-trait correlations, HBA1c, GGT, CRP, ALT and HDL were significantly correlated to LRS. The correlations were all positive except HDL (−0.16 ± 0.04). In females, LRS was significantly correlated to GGT, CRP, ALT, HDL and IGF1, with the correlation with HDL (−0.11 ± 0.03) and IGF1 (−0.09 ± 0.03) being negative.

**Table 2.**
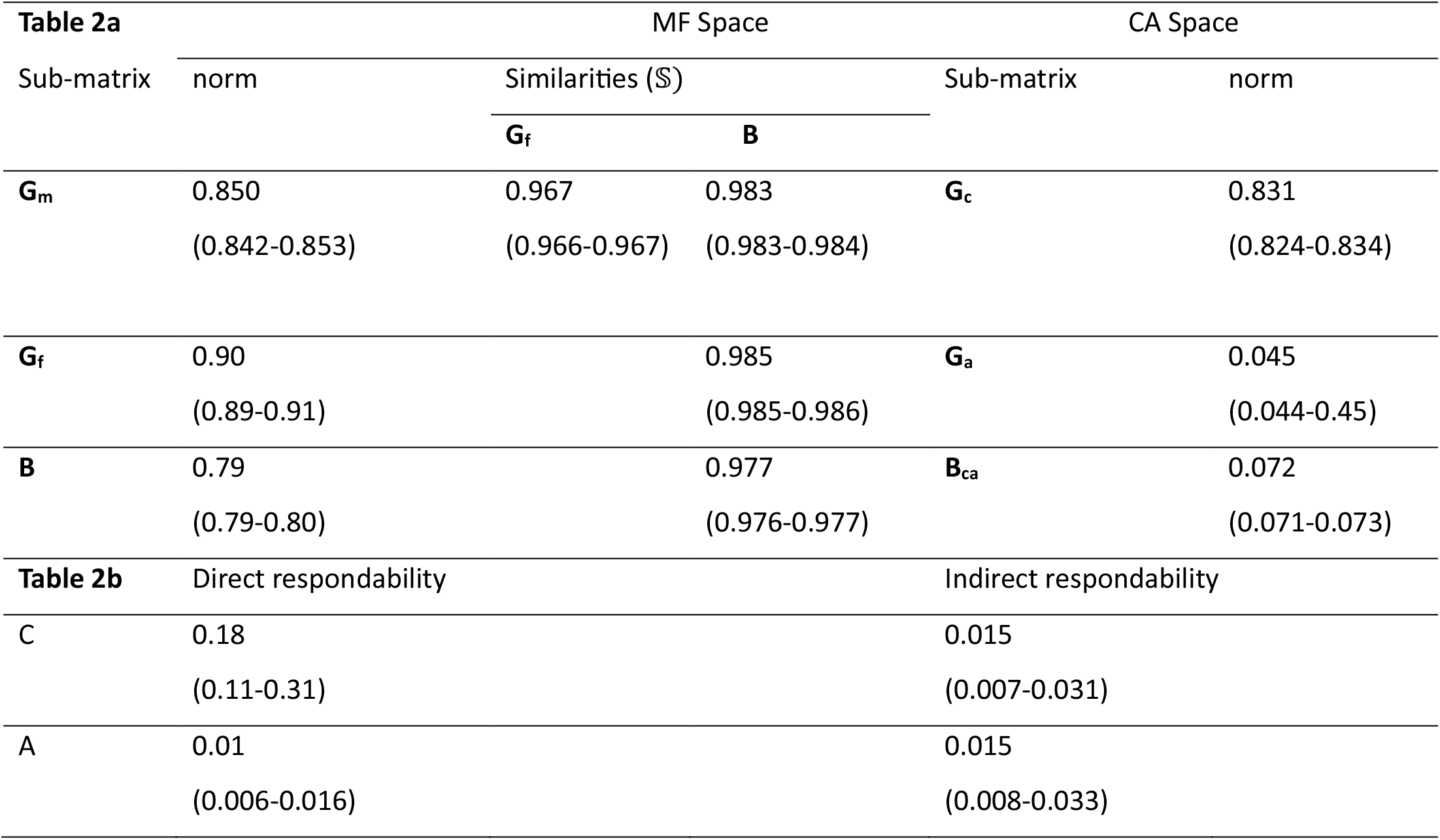
Characteristics of metabolic matrices in male/female (MF) space and concordant/antagonistic (CA) space, and direct and indirect respondabilities. Table 2a depicts the norm of the sub-matrices in MF and CA space as well as the similarities. Values shown are point estimates from the original matrices with 2.5 % and 97.5 % quantile jackknife intervals calculated from 200 LDSC leave-one-out jackknife samples. Table 2b contains the direct and indirect respondabilities to simulated 1000 concordant (C) and antagonistic (A) random skewers. Values shown are the median and 2.5 % and 97.5 % quantile intervals.

**Figure 1:**
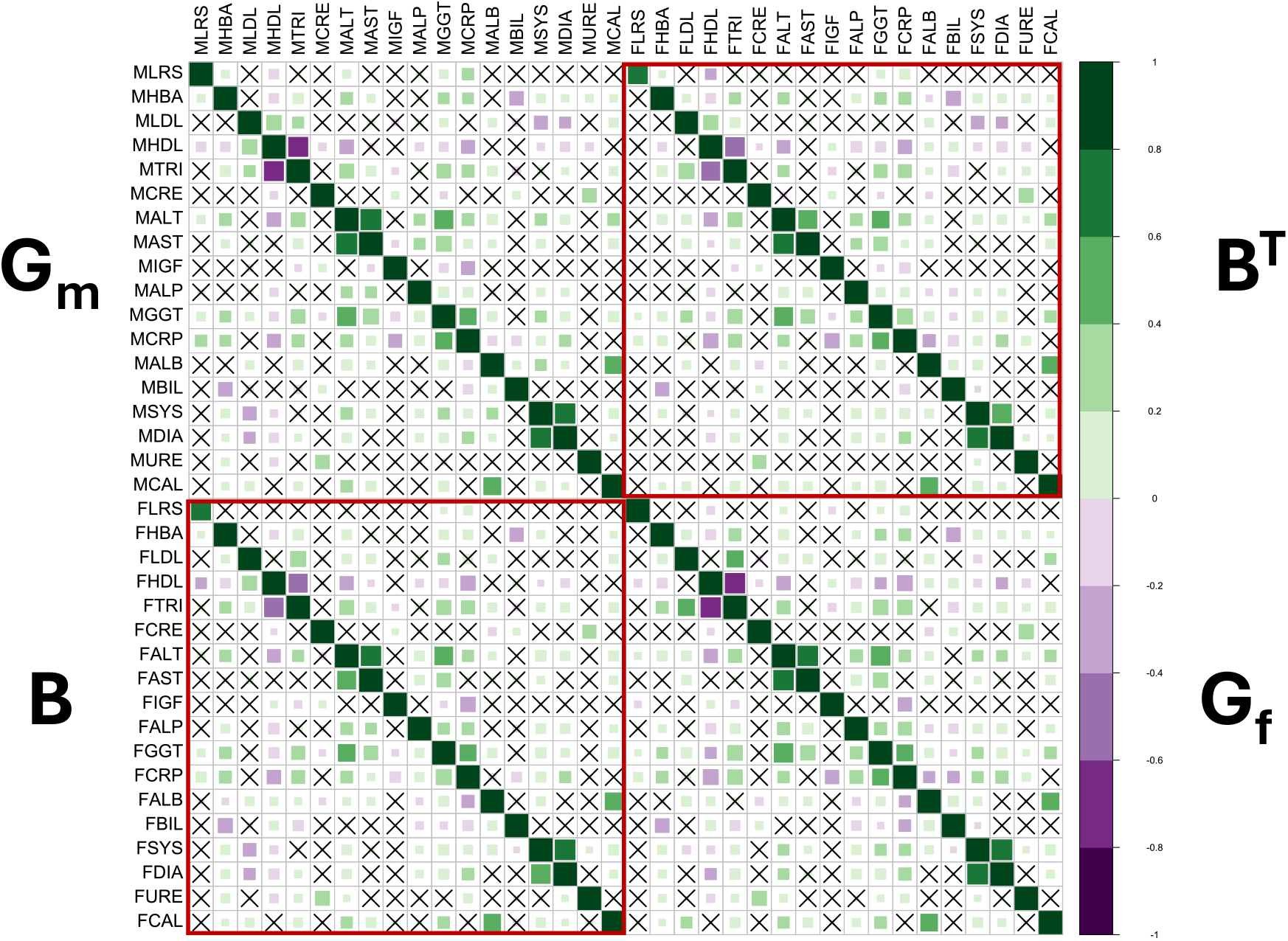
G_mf_ correlation matrix. 36 × 36 genetic correlation matrix of 18 male and female metabolic and physiological traits in humans including lifetime reproductive success (LRS) as a fitness-proxy. The top-left is the male correlation matrix (G_m_), the bottom-right is the female correlation matrix (G_f_), bottom-left is the between-sex genetic correlation matrix (B), and the top-right is the transpose of the B matrix. B and B^T^ are outlined in red. The shades of green represent positive correlation whereas shades of purple represent negative correlations. The size and intensity of colour of the squares represent the strength of correlations. Squares marked with crosses are not significant at FDR <0.05.

The cross-sex genetic correlation or the intersexual genetic correlation for fitness (LRS) in this dataset was 0.61 ± 0.05 and was significantly different from 1 (P = 6.19 × 10^−15^). The cross-sex correlations were overall very high, around ~0.90 for all the traits except LRS (**Supplementary Figure 2, Supplementary Table 4**). The strongest cross-sex genetic correlation was for HBA1c (0.96 ± 0.02) and CRP (0.96 ± 0.02).

### b) G_**m**_ and **G**_**f**_ matrices

We estimated the full genetic (co)variance matrices (**G**_**mf**_) with LDSC regression (**Supplementary Tables 5&6**). To investigate the multivariate genetic architecture in males and females, we did an eigen decomposition of **G**_**m**_ and **G**_**f**_ (covariance matrices). Both were positive-definite and full-rank matrices.

Eigenanalysis revealed that the major eigenvector (**g**_**max**_) of **G**_**m**_ and **G**_**f**_ explained 20% and 22% of genetic variance in males and females, respectively (**Supplementary Table 7&8**) The first 12 out of 18 eigenvectors cumulatively explained 91% and 90% of the total genetic variance in **G**_**m**_ and **G**_**f**_, respectively, indicating that genetic variance was evenly distributed in the phenotypic space and the **G** matrices were not ill-conditioned.

### b) Asymmetry in B

We decomposed **B** into symmetric and skew-symmetric components and estimated the proportion of sum of squares in **B** which is skew-symmetric as a quantification of asymmetry. We found that **B** is only 1.15% asymmetric compared to a null of 0% asymmetry (**Supplementary Table 9**). This indicates that the shared multivariate genetic architecture of males and females for this set of metabolic traits is largely similar. This contrasts with our earlier reported asymmetry in anthropometric and sex-hormonal traits in humans, which was 14.4% [13] highlighting that the degree of asymmetry in **B** is trait-dependent. Hence, for these metabolic traits sex-biased evolutionary response would be limited.

### c) Genetic variation in the concordant-antagonistic subspace and respondabilities

We transformed the male-female space in the **G**_**mf**_ matrix to concordant and antagonistic submatrices using the multivariate framework derived by Cheng and Houle (2020) [19] as described in the Methods section.

We found that the Frobenius norm of **B** is as large as that of **G**_**m**_ and **G**_**f**_ (‖***B***‖ = 0.79, ‖***G***_***m***_‖ = 0.85, ‖***G***_***f***_‖ = 0.94) (**Table 2a, Figure 2A**) with a high similarity of (S = 0.98) between the within-sex covariance matrices and **B**. This is expected given the *r*_*mf*_ of the traits involved are all very high, approaching 1. The similarity between **G**_**m**_ and **G**_**f**_ is also high (S = 0.96) indicating shared genetic variation between the sexes. **B** itself is highly symmetrical as its correlation with **B**^**T**^ is (S = 0.977) which corroborates with our earlier analysis about the percentage of asymmetric components of **B**.

**Figure 2:**
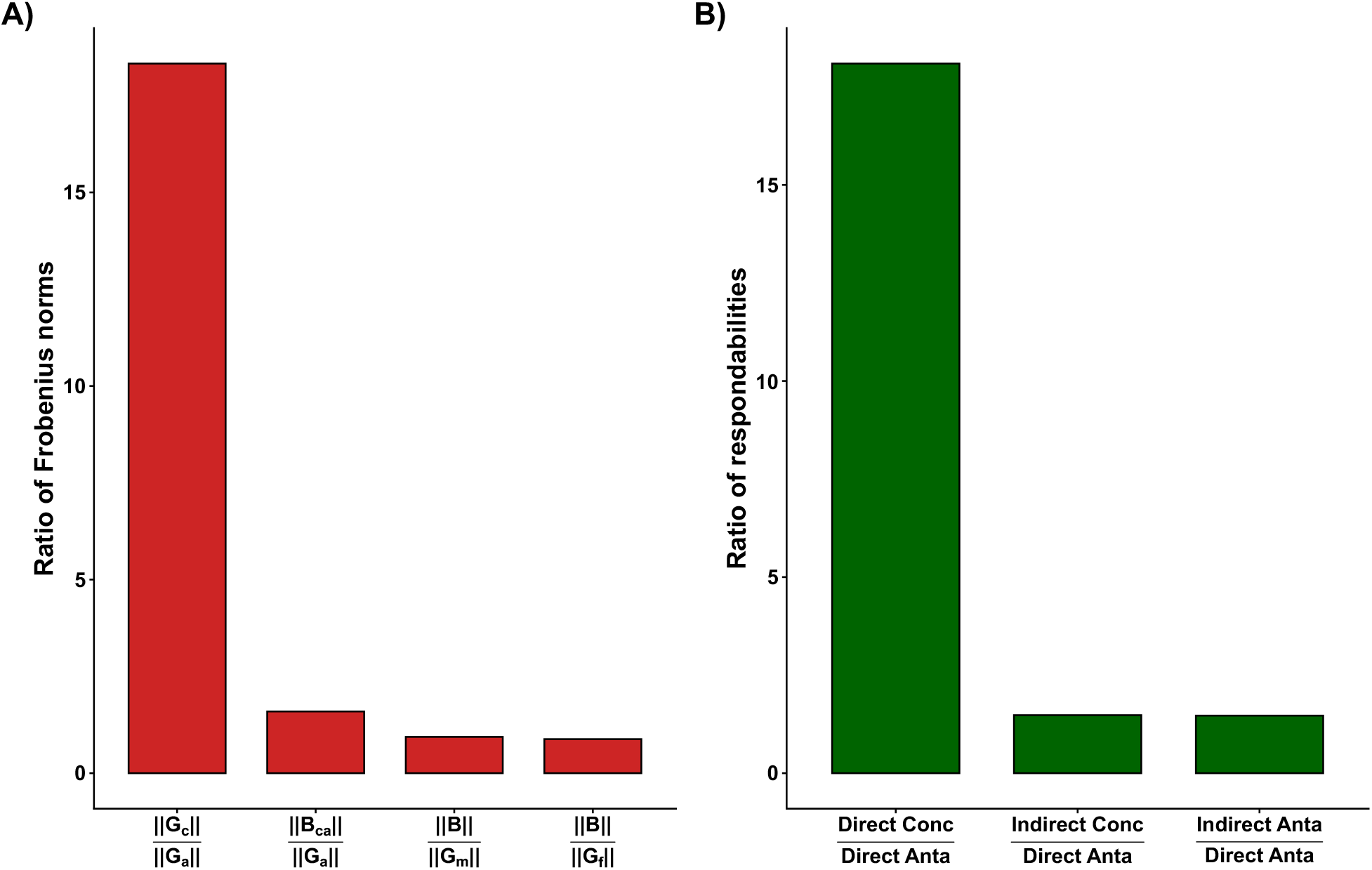
Frobenius norms of concordant and antagonistic subspaces and the respondabilities to concordant and antagonistic selection. A) Ratio of Frobenius norms of **G**_**c**_ and **B**_**ca**_ to **G**_**a**,_ and that of **B** to **G**_**m**_ and **G**_**f**_ indicating that the size of the concordant subspace is 18.5 times as much as the antagonistic subspace. The covariation between concordant and antagonistic subspace (**B**_**ca**_) is 1.6 times larger than G_a_. B) Ratio of respondabilities showing direct concordant evolutionary response exceeds the direct antagonistic response by 18-fold while the indirect concordant response is about 1.4 times that of the direct antagonistic response. ‘Conc’ stands for concordant and ‘Anta’ for antagonistic respondabilities.

In the concordant-antagonistic space, we found that the Frobenius norm of the matrix of concordant genetic variation, **G**_**c**_, is about 18.5-fold larger than that of the size of the antagonistic matrix **G**_**a**_ (‖***G***_***c***_‖ = 0.831, ‖***G***_***a***_‖ = 0.045) (**Figure 2A**) which suggests that the genetic architecture facilitates evolutionary response far more strongly along concordant than antagonistic dimensions. On the other hand, the norm of the matrix of genetic covariation between antagonistic and concordant subspaces, **B**_**ca**_ is about 1.6-fold larger than that of the norm of **G**_**a**_, (**Figure 2A**) indicating larger potential for indirect responses to concordant selection compared to direct antagonistic responses.

Consistent with this, we found that the direct respondability, i.e., the predicted responses to concordant selection is much larger than direct responses to antagonistic selection (**Table 2b**). Interestingly, the indirect respondability to concordant selection is about 1.5 times larger than that of direct respondability to antagonistic selection (**Figure 2B**), suggesting sex differences in these metabolic traits is more likely to evolve via indirect response to concordant selection compared to direct response to antagonistic selection.

The quantile intervals for the matrix norms are derived from LDSC leave-one-block-out jackknife samples. As these pseudo-estimates differ only slightly from the full-data estimates, the resulting intervals were narrow.

### d) Role of B in the evolution of sexual dimorphism

To evaluate the evolutionary consequences of cross-sex genetic covariance, we simulated SC and SA random selection vectors and used the multivariate breeder’s equation to find the predicted short-term response to selection in the sexes, once constraining **B** to zero and once including it. A significant reduction in the angle between male and female response (**Δz**_**m**_ and **Δz**_**f**_) while including vs excluding **B** is an indication that **B** acts as a constraint to the evolution of sexual dimorphism by reducing the angle and forcing males and females to evolve in similar direction.

When we simulated SC selection, the median angle between **Δz**_**m**_ and **Δz**_**f**_ without **B** was observed to be small, 13° (median) [2.5% and 97.5% quantile intervals: 8.6° - 23°]. Including **B**, this was further reduced to 9° [5.5° - 13.5°] (**Figure 3**). Because the quantile intervals overlapped, we concluded that **B** neither facilitates nor constrains the evolution of sexual dimorphism under concordant selection. Under completely SA selection (**θ** between **β**_**m**_ and **β**_**f**_ = 180°), which is an extreme case, this angle, before the inclusion of **B** was large and in virtually opposite directions, 167° [157° - 171.4 °], and with **B** it reduced to 65.9° [30.45° - 108°]. This was a significant reduction of the angle, about 65%, which indicates that **B** acts as a strong constraint to the evolution of sex differences under antagonistic selection, and a symmetric **B** adds to this constraint.

**Figure 3:**
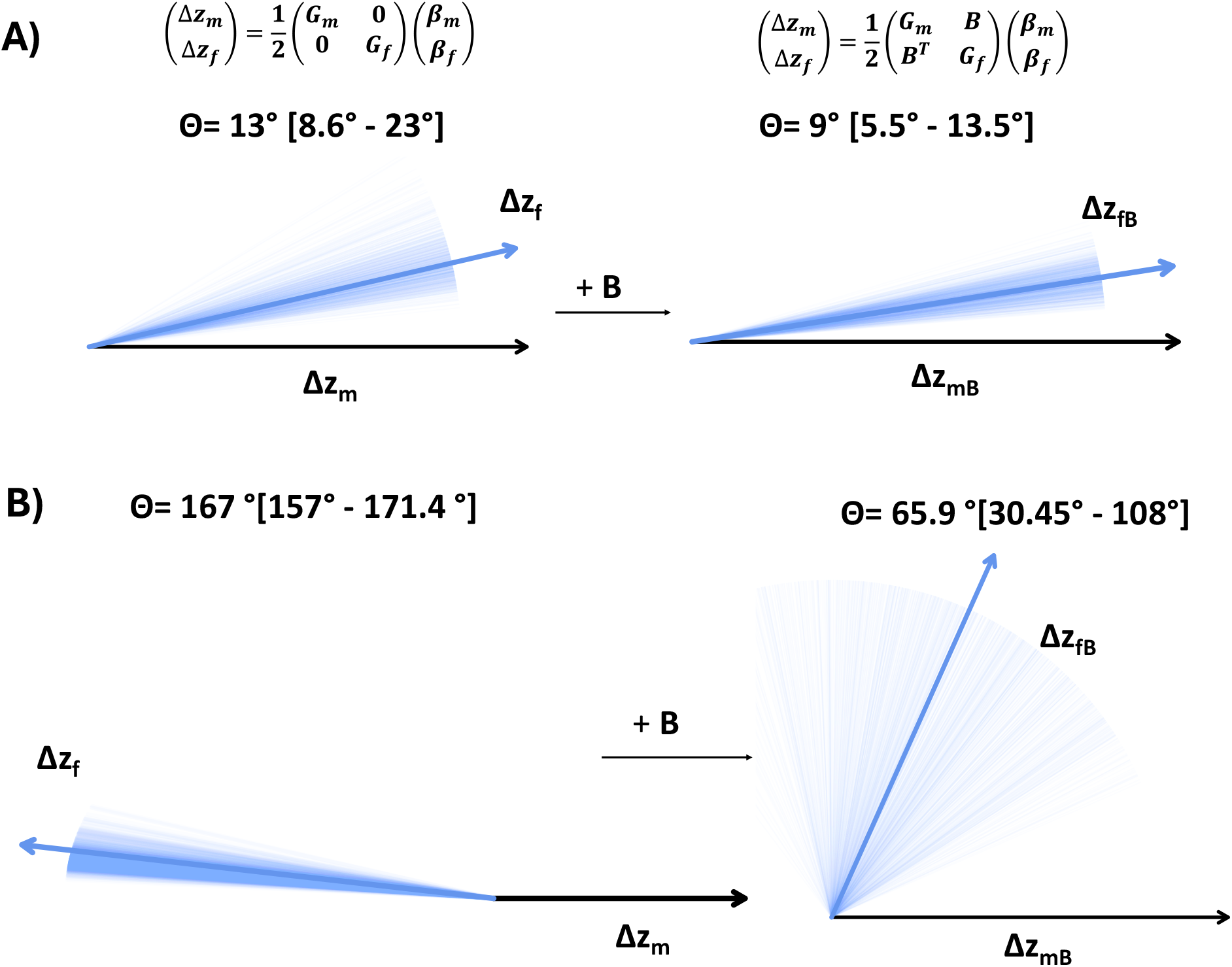
Evolutionary simulation with random selection vectors using multivariate breeder’s equation. A) Median Angles between predicted short-term response in males (**Δz**_**m**_) and females (**Δz**_**f**_) to 1000 random sexually concordant selection vectors with **B** constrained to zero, and after including **B** (**Δz**_**mB**_ and **Δz**_**fB**_). Thin lines represent the 1000 response vectors, and the thick arrow is the median response. B) Median angle between male and female response vectors when selection is sexually antagonistic. **B** constraints the evolution of sexual dimorphism in this case by reducing the angle between male and female responses by 65%. The length of the vectors is only for representation.

Together, the small degree of asymmetry and genetic constraint of **B** indicate males and females have a largely similar multivariate genetic architecture which might impede evolutionary divergence. Indeed, our analysis showed that **B** primarily acts as a constraint for metabolic traits under SA selection rather than SC selection. However, our previous disentanglement of concordant and antagonistic subspaces revealed that there is very little antagonistic genetic variation for SA selection to act upon with indirect responses to SC selection being stronger than direct response to SA selection.

### e) Local genetic correlations and concordant and antagonistic regions

We next examined local SNP-heritabilities, cross-sex genetic correlations, and cross-sex-cross-trait genetic correlations across 1691 independent genomic regions to assess evidence for IASC. Analyses were restricted to traits that showed significant genome-wide genetic correlations with LRS in at least one sex. These included five traits in males, HbA1c, GGT, CRP, ALT, and HDL, and five in females, GGT, CRP, ALT, HDL, and IGF1, with a union of six traits (**Supplementary Figure 4**).

Across these traits, we estimated local SNP-heritabilities and genetic correlations for 21 cross-sex-cross-trait combinations and 6 cross-sex single trait combinations (**Figure 4**; **Supplementary Figures 5-9, Supplementary Tables 17-60**). We found significantly correlated local regions, both positive and negative, throughout the genome for all the cross-sex-cross-trait genetic correlations at a Bonferroni adjusted threshold of P < 2.96 × 10^−5^ for each trait pair. In contrast, all local cross-sex genetic correlations for the same trait measured in males and females were positive, indicating an absence of local antagonism when traits are considered in isolation.

**Figure 4:**
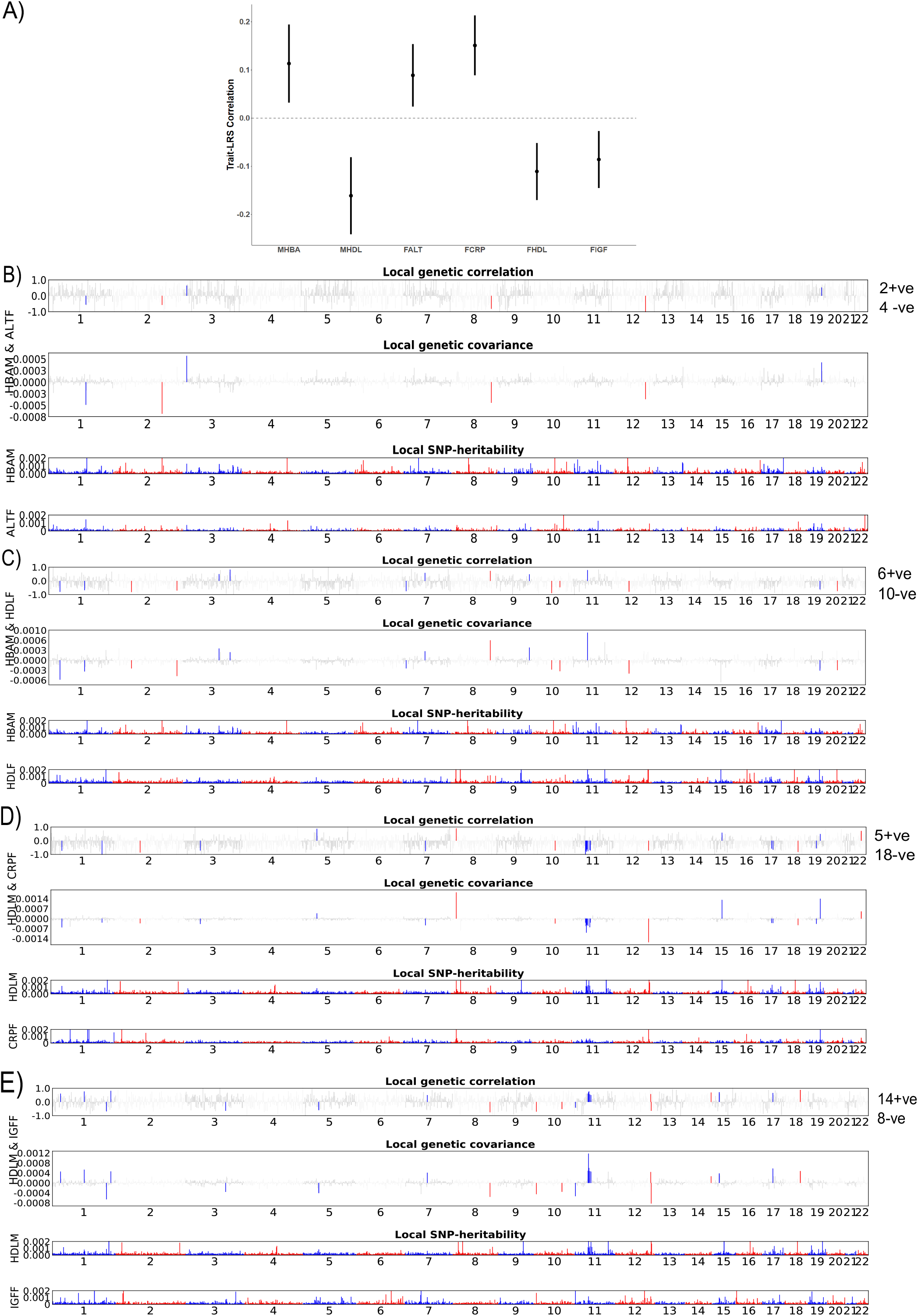
Local cross-sex-cross-trait genetic correlations and antagonistic and concordant regions. A) Trait-fitness genetic correlation for male HbA1c (MHBA), male HDL (MHDL), female ALT (FALT), female CRP (FCRP), female HDL (FHDL), female IGF1 (FIGF). B) Manhattan-style plot of local genetic correlation and covariance between MHBA and FALT. X-axis contains the chromosomes from 1-22, and along the Y-axis are the genetic covariances/correlations. The significant regions are marked in colour, red for even and blue for odd chromosomes. Lower panel contains the local SNP-heritability of the two traits. Both MHBA and FALT are positively correlated to LRS; hence the local positively correlated regions are concordant while the negatively correlated regions are antagonistic. C) Local correlations and covariance between MHBA and FHDL. MHBA is positively correlated to LRS while FHDL is negatively. Therefore, the positively correlated regions are antagonistic while the negatively correlated regions are concordant. D) Local correlations and covariance between MHDL and CRPF. MHDL is negatively correlated to LRS and CRP is positively. The positively correlated local genomic regions are antagonistic while negative regions are concordant. E) Local correlations and covariances between MHDL and FIGF, both of which are negatively correlated to LRS. Positively correlated local regions are concordant while negatively correlated regions are antagonistic.

Notably, 2 out of 24 trait combinations did not show significant genome-wide cross-sex-cross-trait genetic correlations nevertheless exhibited significant local correlations. For example, male HbA1c-female IGF1 (genome-wide r_g_ = −0.005 ± 0.03, P_adj_ = 0.87) and male ALT-female IGF1 (r_g_ = 0.007 ± 0.03, P = 0.86) displayed 7 and 8 genomic regions, respectively, with significant local correlations (**Supplementary Figures 9 & 6)**. These findings highlight the importance of local analyses for detecting correlated genomic regions that is masked or averaged out at the genome-wide level.

To interpret local correlations in the context of IASC, we classified genomic regions as concordant, antagonistic, or mixed based on the sign of the local cross-sex-cross-trait genetic correlation relative to the direction of trait-LRS associations (see **Table 1** in Methods). Concordant regions were defined as those in which local cross-sex-cross-trait genetic effects were aligned with fitness in both sexes, whereas antagonistic regions showed opposite fitness directions between males and females.

We identified 138 significant genomic regions across all cross-sex-cross-trait combinations. Of these, 68% were consistently concordant, 12% were consistently antagonistic, and the remaining 20% showed mixed effects (**Supplementary Table 10**), being concordant in some trait combinations and antagonistic in others.

We found that genome-wide concordance can hide significant local conflict. Among 10 out of 24 trait pairs with significant positive genome-wide cross-sex-cross-trait genetic correlations exhibited at least one genomic region with significant negative local correlation. For example, male HbA1c-female ALT (r_g_= 0.22 ± 0.04) and male GGT-female CRP (0.40 ± 0.07) both showed positive global correlations, reflecting shared genetic influences that increase both traits and are also positively associated with LRS in both the sexes. Despite this genome-wide concordance, we detected multiple genomic regions with significant negative local correlations for these same trait pairs (four regions for male HbA1c-female ALT and two regions for male GGT-female CRP) (**Figure 3 & Supplementary Figure 7**). In these regions, alleles that increase the trait in one sex decrease the correlated trait in the other sex, thereby generating opposite fitness consequences across sexes.

The local correlations of trait pairs where the correlation with LRS is opposite in the sexes are particularly informative for understanding IASC. For example, male HBA1c and female HDL are positively and negatively correlated to LRS, respectively (**Supplementary Figure 4**) and their global genetic correlation is negative (Corr (male HBA1c, female HDL) = −0.16 ± 0.03). Their local correlation shows both significant positive (seven) and negative (ten) regions (**Figure 3**). In regions with positive local genetic correlation, alleles increase both traits in males and females. However, because HbA1c and HDL have opposite fitness associations in the two sexes, such alleles generate opposing fitness effects, benefiting one sex while causing harm in the other, and therefore represent locally antagonistic regions. In contrast, regions with negative local genetic correlation exhibit opposing allelic effects on the two traits; in this case, these effects lead to reduced fitness in both sexes, resulting in local concordance rather than conflict. This contrasts with the previous case where the negatively correlated regions were antagonistic. Thus, it is the opposing allelic effects on fitness rather than the sign of the cross-sex-cross-trait correlation that determines whether a region is under IASC. Thus, local antagonism and conflict can persist even when genome-wide genetic correlations are concordant. Genome-wide approaches may therefore underestimate the extent and genomic distribution of IASC.

Among recurrently significant regions, we identified three independent regions that were predominantly concordant (chr12:119,754,110-122,007,651 bp; 9/9 concordant), predominantly antagonistic (chr22:37,570,269-39,307,894 bp; 5/7 antagonistic), and mixed (chr7:71,874,885-73,334,602 bp; 7 concordant and 6 antagonistic; **Supplementary Table 11**). Patterns of SNP-heritability across these regions further suggested heterogeneity in the genetic basis of local conflict. In predominantly concordant regions, 9/9 trait combinations showed significant local SNP-heritability, whereas in predominantly antagonistic regions, 3/5 trait combinations exhibited significant heritability in only one of the two traits (**Supplementary Table 11**). This asymmetry suggests that antagonistic regions may be characterized by sex-limited genetic effects, where genetic variation influences phenotype primarily in one sex, whereas concordant regions involve genetic variation with effects in both sexes.

These results demonstrate that IASC in metabolic traits is genomically localized and multivariate in nature. Importantly, global genetic correlations, whether concordant or antagonistic, conceal the extensive heterogeneity in local genetic architecture, with individual regions harbouring alleles with concordant, antagonistic, or mixed fitness consequences.

### f) Gene mapping and pathway analyses

To gain biological insights into the mechanisms underlying conflict, we functionally annotated the genomic regions as previously classified – predominantly concordant, predominantly antagonistic, and mixed. We hypothesized that these regions would exhibit distinct functional profiles reflecting their roles in sex-specific fitness architecture. Because local regions often differed in SNP-heritability across traits, we selected, for each region type, the trait with the highest local SNP-heritability as the representative phenotype for gene mapping. Male CRP had the highest local heritability in the concordant region (h^2^_local_ = 0.00119 ± 0.00048), female CRP in the antagonistic region (h^2^_local_ = 0.0008± 0.00015), and female GGT in the mixed region (h^2^_local_ = 0.0017 ± 0.000165) (**Supplementary Table 12**). We mapped genes using the GWAS summary statistics for these traits in those specific regions.

Total 53 genes were mapped to male CRP (**Supplementary Table 13**), 33 genes to female CR**P** (**Supplementary Table 14**), and 27 genes to female GGT (**Supplementary Table 15**). Functional enrichment analysis revealed that the predominantly concordant region was significantly enriched for pathways related to synaptic transmission (P_adj_ =0.02; Elsevier Pathways) and nominally significant for elevation of cytosolic calcium ion levels (P = 0.00033; unadjusted; Reactome Pathways) (**Supplementary Table 16**). Genes in the predominantly antagonistic genomic region showed enrichment for canonical WNT signalling pathway across multiple annotation databases with adjusted P-value significant in one (P_adj_ = 0.048; Elsevier Pathways). In addition, the genes from the latter region were also nominally significant for ovarian infertility in (P = 0.04, unadjusted; WikiPathways Mouse) and meiosis (P=0.009, unadjusted; Reactome Pathways), suggesting possible links between antagonistic genetic effects and reproductive or developmental processes. On the other hand, the mixed region genes were enriched for 7q11.23 copy number variation locus (P_adj_ = 1.37 × 10^−19^; WikiPathways) which is a highly pleiotropic region associated with Williams-Beuren syndrome [57], and DNA repair pathways (P_adj_ = 0.03; MSigDB Hallmark). Overall, these results indicate that genomic regions associated with concordant, antagonistic, and mixed patterns of local genetic correlation are characterized by partially distinct functional signatures alluding to different biological processes associated with these categories. Since most of the pathways were enriched by not more than two genes in the gene-set (except 7q11.23 CNV locus – 10 genes), we take these results as suggestive.

## Discussion

In this study, we showed that the multivariate genetic architecture of metabolic traits in humans is largely shared between the sexes with the presence of local sexually antagonistic (SA) genetic variation. The amount of multivariate genome-wide sexually concordant (SC) variation is much larger than that of SA variation with indirect responses to SC selection being stronger than direct responses to SA selection. The **B** matrix was largely symmetric and constrained the evolution of sex difference under random SA and not SC selection. The genetic associations of the traits with LRS were of the same sign in both the sexes highlighting the lack of antagonistic trait-fitness correlation. Given the sexual dimorphism in these traits, one of the possible explanations for the between-sex shared architecture is a weak or largely resolved genome-wide conflict where the sex differences might arise from sex-specific regulatory mechanisms. In contrast, local genetic correlation analyses uncovered a relatively small but non-negligible proportion of genomic regions under consistent conflict, with the majority showing concordant effects. Importantly, these different classes of regions were enriched for distinct biological pathways, highlighting that genome-wide summaries can hide biologically meaningful signatures of intralocus sexual conflict (IASC) at finer genomic scales.

### a) A symmetric B matrix and the predominance of concordant genetic variation at the genome-wide level

The canonical model of the evolution of sexual dimorphism largely relies on SA selection acting on homologous traits in males and females and the genetic constraint faced due to a shared genetic architecture. However, dimorphism can evolve in the absence of antagonistic selection (a) when selection differs in magnitude, or (b) there is difference in genetic variances for homologous traits between the sexes. Cheng and Houle (2020) [19] analytically showed that SC selection can be as important as SA selection in the evolution of dimorphism, depending on the genetic architecture. We applied this framework in human metabolic traits and found that the amount of SC genetic variation is 18.5-fold larger than SA variation. Moreover, the amount of genetic covariation between antagonistic and concordant subspaces is 1.6-fold larger than SA variation itself. Furthermore, in concordance with the above finding, we showed that indirect responses to SC selection is much stronger than direct responses to SA selection. The predominance of SC genetic variation and higher indirect respondability to SC selection is expected given the high cross-sex correlations and an almost symmetric **B** matrix. Thus, despite the existence of local antagonistic regions, the global multivariate genetic architecture is dominated by concordant genetic variation, causing evolutionary responses to be influenced primarily by concordant rather than antagonistic subspaces. A meta-analyses testing the Cheng and Houle framework on published **B** matrices across several traits and species revealed that the overall amount of SC genetic variation is much higher than SA variation and implied the role of indirect responses to SC as a potential mechanism of evolution of sex differences [17]. Our results suggest that, given the observed genetic architecture, sexual dimorphism in metabolic traits may evolve primarily through indirect responses to SC selection, with sex differences plausibly arising as correlated responses to selection on other traits rather than through strong SA selection on the focal traits themselves.

Our study showed that the **B** matrix representing the shared genetic architecture for a set of metabolic and physiological traits in humans is largely symmetric between the sexes indicating that the cross-sex-cross-trait covariances are not different based on the sex where they are expressed. A recent study has shown that across several taxa, **B** has a wide range of asymmetry (0.005 to 0.78) with little dependence on trait types or species [17]. The only other asymmetry information available in humans is that of anthropometric and sex-hormonal traits showing moderate estimate (0.14) [13]. These limited data suggest that **B** matrix asymmetry in humans may be lower than the cross-taxa average, and potentially trait-dependent, which calls for further investigation. We found that the symmetric **B** acts as a constraint to the evolution of dimorphism under random SA selection but not SC selection. The absence of antagonistic trait-fitness correlations suggests that, for metabolic traits, IASC is not currently driven by strong genome-wide SA selection. Since these traits are phenotypically sexually dimorphic [58–60], we surmise two possible explanations for the observed results– (a) global or genome-wide IASC is weak and might have been resolved through sex-specific regulation with the evolution of dimorphism and a subsequent increase in r_mf_s and reduction of asymmetry in **B**. (b) Alternatively, strong genome-wide conflict never existed for this set of metabolic traits and the sex differences primarily evolved as an indirect response to concordant selection. The r_mf_ as well as the structure of **B** after the evolution of dimorphism depends on the strength of selection, mutation rate and the evolved genetic architecture [1]. Currently, we have little knowledge about the dynamics of **B** in IASC both theoretically and empirically [5, 10], but our study highlights the consequences of a symmetric **B**. This study adds an important dimension to the empirical body of work showing the persistence of sexual dimorphism despite a lack of sex-specific multivariate genetic architecture in metabolic traits in humans.

### b) Local genetic architecture shows a mosaic of concordant and antagonistic genomic regions

Our local analyses unfolded a more nuanced picture. Consistent with the genome-wide analyses, the local cross-sex-cross-trait genetic correlations revealed that most of the regions were concordant, i.e., their association with genome-wide fitness were in the same direction in both the sexes, indicating concordance predominates at both genome-wide and local scales. However, a small number of genomic regions were consistently antagonistic and represented regions of persistent local conflict. Interestingly, a third category of regions emerged which were under conflict for some trait combinations whereas concordant for others. These genomic regions may reflect constraints arising from correlated traits due to pleiotropy or LD, such that the same genes are under ongoing conflict for a trait pair whereas represent resolved conflict for other trait combinations. Moreover, these local correlations illustrate that despite a similar genome-wide correlation, the local genetic architecture can be distinctly different between traits across sexes. However, we note there are at least two limitations of the scheme of classification we have used to define regions under conflict – a) using LRS as a single fitness proxy which is debatable in contemporary humans, but is widely used [28] and b) using genome-wide trait-fitness correlations to link fitness outcomes at the local level. We must also highlight that while ideally local trait-LRS associations would directly test local fitness effects, power limitations prevented detection of significant SNP-heritability in local regions for LRS in both the sexes, which might have happened due to low genetic variance in those regions which did not survive multiple testing correction. Moreover, our focus on significantly correlated local regions ensured we identify only the strongest signals, meaning our estimates of antagonistic and concordant regions are likely conservative lower bounds. The true extent of sexually antagonistic and concordant architecture across the genome is probably substantially larger, involving many smaller-effect loci that fall below our detection threshold.

### c) Functional enrichment of local regions

We found that our categorisation of antagonistic and concordant regions were not mere statistical artifacts but show suggestively distinct functional enrichment, though modest in magnitude. The predominantly concordant region was enriched for synaptic transmission and nominally for calcium signalling pathways. Neuronal calcium signalling has been linked to metabolic regulation in *Drosophila [61]* and calcium homeostasis to lipid metabolism [62]. Moreover, sex differences in calcium signalling and homeostasis have been reported [63, 64] and dysregulation of calcium signalling might be a key aspect of cardiovascular pathogenesis. These pathways indicate biological functions that have sex-specific regulatory mechanisms and may contribute to resolution of conflict. On the other hand, the predominantly antagonistic region was enriched for canonical WNT signalling pathway and nominally significant for meiosis, and ovarian infertility in mouse. Canonical WNT signalling or β-catenin dependent pathway is highly conserved across taxa, suppression of which is involved in embryonic testis development while activation leads to development of ovaries [65, 66]. In fact, it also plays important role in adult male and female reproductive physiology and pathogenesis [67–70]. Moreover, the activation of WNT signalling pathway is associated with sexual dimorphism observed in non-alcoholic fatty liver disease (NAFLD) [71], and the metabolic traits in our study consist of a subset of the biomarkers for this disease involving liver enzymes, bilirubin, albumin, creatinine, CRP, etc., which might explain the enrichment of this specific pathway. These biological features, combined with opposite fitness effects between sexes, are consistent with ongoing local ontogenic sexual conflict mediated through the pleiotropic metabolic and developmental pathways such as WNT signalling.

The metabolic and physiological traits involved in this study are themselves sexually dimorphic and are disease biomarkers for disorders showing sex difference. But our study revealed a largely shared predominantly concordant genome-wide genetic architecture between the sexes with antagonism in local regions. This implies that the sex difference in these traits arises due to sex-specific regulation of a shared genetic architecture [72–74]. Sex-biased gene expression [75–77], sex-specific alternative splicing [78], and epigenetic regulation [79, 80], etc. can give rise to sex differences in absence of sex-specific genetic variants or sex-specific effect sizes of the same variants, through downstream differential regulation. These are also some of the mechanisms of resolution of IASC [1] and the current genome-wide shared genetic architecture between sexes hints towards a largely resolved past conflict or absence of strong conflict, with ongoing conflict at the local level. However, further work integrating regulation, gene-expression and epigenetics are required to uncover the underlying sex-specific mechanisms involved in the evolution of sex differences in these metabolic traits.

In summary, our study showed a shared between-sex multivariate genetic architecture of human metabolic traits which are sexually dimorphic phenotypes along with a preponderance of concordant genetic variation. We observed that the indirect responses to concordant selection is stronger than direct responses to antagonistic selection. However, there were local antagonistic regions representing ongoing conflict, albeit in much smaller scale. This mosaic of antagonism and concordance at the local level suggests that IASC resolution is neither complete nor uniform across the genome, but rather an ongoing evolutionary process with different genomic regions at different stages of resolution. These results highlights the importance of concordant genetic variation at the genome-wide as well as local levels and underscores its role in sex differences in humans. Our work is a significant contribution to the study of IASC in humans and emphasizes the necessity and importance of integrating multivariate and local genetic architecture to understand the biology and evolution of sexual conflict.

## Acknowledgements

AC was funded by INSPIRE Faculty Fellowship, Department of Science and Technology, Govt. of India, with registration number, IFA20-LSBM-241.

## Data accessibility

We used publicly available GWAS summary statistics for all the analyses, downloaded from Neale lab webpage (https://www.nealelab.is/uk-biobank). Representative codes for estimating genome-wide and local genetic correlations using LDSC and ρ-HESS, respectively, are given in the supplementary material.

## References

[1] Bonduriansky, R. & Chenoweth, S. F. 2009 Intralocus sexual conflict. Trends Ecol. Evol. 24, 280–288.

[2] Van Doorn, G. S. 2009 Intralocus Sexual Conflict. Ann. N. Y. Acad. Sci. 1168, 52–71. (DOI:10.1111/j.1749-6632.2009.04573.x).

[3] Long, T. A. F. & Rice, W. R. 2007 Adult locomotory activity mediates intralocus sexual conflict in a laboratory-adapted population of Drosophila melanogaster. Proc. R. Soc. B 274, 3105–3112. (DOI:10.1098/rspb.2007.1140).

[4] van der Bijl, W. & Mank, J. E. 2021 Widespread cryptic variation in genetic architecture between the sexes. Evolution Letters 5, 359–369. (DOI:10.1002/evl3.245).

[5] Pennell, T. M., Mank, J. E., Alonzo, S. H. & Hosken, D. J. 2024 On the resolution of sexual conflict over shared traits. Proc. R. Soc. B 291. (DOI:10.1098/rspb.2024.0438).

[6] Lande, R. 1980 Sexual dimorphism, sexual selection, and adaptation in polygenic characters. Evolution 34, 292–305.

[7] Poissant, J., Wilson, A. J. & Coltman, D. W. 2010 Sex-specific genetic variance and the evolution of sexual dimorphism: a systematic review of cross-sex genetic correlations. Evolution 64, 97–107.

[8] Puixeu, G. & Hayward, L. K. 2025 The relationship between sexual dimorphism and intersex correlation: do models support intuition? Genetics 231. (DOI:10.1093/genetics/iyaf175).

[9] Lande, R. 1979 Quantitative genetic analysis of multivariate evolution, applied to brain: body size allometry. Evolution 33, 402–416.

[10] McGlothlin, J. W., Cox, R. M. & Brodie III, E. D. 2019 Sex-specific selection and the evolution of between-sex genetic covariance. J. Hered. 110, 422–432.

[11] Gosden, T. P. & Chenoweth, S. F. 2014 The evolutionary stability of cross-sex, cross-trait genetic covariances. Evolution 68, 1687–1697.

[12] Ingleby, F., Innocenti, P., Rundle, H. & Morrow, E. 2014 Between-sex genetic covariance constrains the evolution of sexual dimorphism in Drosophila melanogaster. J. Evol. Biol. 27, 1721–1732.

[13] Chakrabarty, A., Chakraborty, S., Nandi, D. & Basu, A. 2024 Multivariate genetic architecture reveals testosterone-driven sexual antagonism in contemporary humans. PNAS 121, e2404364121. (DOI:doi:10.1073/pnas.2404364121).

[14] Delph, L. F., Andicoechea, J., Steven, J. C., Herlihy, C. R., Scarpino, S. V. & Bell, D. L. 2011 Environment-dependent intralocus sexual conflict in a dioecious plant. New Phytol. 192, 542–552. (DOI:10.1111/j.1469-8137.2011.03811.x).

[15] Hawkes, M., Lane, S. M., Rapkin, J., Jensen, K., House Clarissa M., Sakaluk, S. K. & Hunt, J. 2022 Intralocus sexual conflict over optimal nutrient intake and the evolution of sex differences in life span and reproduction. Funct. Ecol. 36, 865–881. (DOI:10.1111/1365-2435.13995).

[16] Iglesias, P. P., Machado, F. A., Llanes, S., Hasson, E. & Soto, E. M. 2023 Opportunities and Constraints Imposed by the G matrix of Drosophila buzzatii Wings. Evolutionary Biology 50, 127–136. (DOI:10.1007/s11692-022-09593-x).

[17] Videlier, M. & Sztepanacz, J. L. 2025 Asymmetry in Cross-Sex Cross-Trait Genetic Covariances and the Evolvability of Sexual Dimorphism. Am Nat 206, 362–374. (DOI:10.1086/737019).

[18] Cox Robert M. & Ryan Calsbeek. 2009 Sexually Antagonistic Selection, Sexual Dimorphism, and the Resolution of Intralocus Sexual Conflict. Am Nat 173, 176–187. (DOI:10.1086/595841).

[19] Cheng, C. & Houle, D. 2020 Predicting Multivariate Responses of Sexual Dimorphism to Direct and Indirect Selection. Am Nat 196, 391–405. (DOI:doi:10.1086/710353).

[20] Morrissey, M. B. 2016 Meta-analysis of magnitudes, differences and variation in evolutionary parameters. J. Evol. Biol. 29, 1882–1904. (DOI:10.1111/jeb.12950).

[21] Sztepanacz, J. L. & Houle, D. 2019 Cross-sex genetic covariances limit the evolvability of wing-shape within and among species of Drosophila. Evolution 73, 1617–1633. (DOI:10.1111/evo.13788).

[22] Houle, D. & Cheng, C. 2021 Predicting the Evolution of Sexual Dimorphism in Gene Expression. Mol. Biol. Evol. 38, 1847–1859. (DOI:10.1093/molbev/msaa329).

[23] Shi, H., Mancuso, N., Spendlove, S. & Pasaniuc, B. 2017 Local Genetic Correlation Gives Insights into the Shared Genetic Architecture of Complex Traits. The American Journal of Human Genetics 101, 737–751. (DOI:10.1016/j.ajhg.2017.09.022).

[24] Werme, J., van der Sluis, S., Posthuma, D. & de Leeuw, C. A. 2022 An integrated framework for local genetic correlation analysis. Nat. Genet. 54, 274–282. (DOI:10.1038/s41588-022-01017-y).

[25] Chippindale, A. K., Gibson, J. R. & Rice, W. R. 2001 Negative genetic correlation for adult fitness between sexes reveals ontogenetic conflict in <i>Drosophila</i>. PNAS 98, 1671–1675. (DOI:doi:10.1073/pnas.98.4.1671).

[26] Stulp, G., Kuijper, B., Buunk, A. P., Pollet, T. V. & Verhulst, S. 2012 Intralocus sexual conflict over human height. Biol. Lett. 8, 976–978. (DOI:10.1098/rsbl.2012.0590).

[27] Archer, C. R., Recker, M., Duffy, E. & Hosken, D. J. 2018 Intralocus sexual conflict can resolve the male-female health-survival paradox. Nature Communications 9, 5048. (DOI:10.1038/s41467-018-07541-y).

[28] Sanjak, J. S., Sidorenko, J., Robinson, M. R., Thornton, K. R. & Visscher, P. M. 2018 Evidence of directional and stabilizing selection in contemporary humans. PNAS 115, 151–156. (DOI:doi:10.1073/pnas.1707227114).

[29] Stearns, S. C., Govindaraju, D. R., Ewbank, D. & Byars, S. G. 2012 Constraints on the coevolution of contemporary human males and females. Proc. R. Soc. B 279, 4836–4844. (DOI:10.1098/rspb.2012.2024).

[30] Cole, J. M., Scott, C. B., Johnson, M. M., Golightly, P. R., Carlson, J., Ming, M. J., Harpak, A. & Kirkpatrick, M. 2024 The battle of the sexes in humans is highly polygenic. PNAS 121, e2412315121. (DOI:doi:10.1073/pnas.2412315121).

[31] Ruzicka, F., Holman, L. & Connallon, T. 2022 Polygenic signals of sex differences in selection in humans from the UK Biobank. PLoS Biol. 20, e3001768. (DOI:10.1371/journal.pbio.3001768).

[32] Lucotte, E. A., Laurent, R., Heyer, E., Ségurel, L. & Toupance, B. 2016 Detection of Allelic Frequency Differences between the Sexes in Humans: A Signature of Sexually Antagonistic Selection. Genome Biology and Evolution 8, 1489–1500. (DOI:10.1093/gbe/evw090).

[33] Zhu, C., Ming, M. J., Cole, J. M., Edge, M. D., Kirkpatrick, M. & Harpak, A. 2023 Amplification is the primary mode of gene-by-sex interaction in complex human traits. Cell Genomics 3. (DOI:10.1016/j.xgen.2023.100297).

[34] Ruzicka, F. & Connallon, T. 2022 An unbiased test reveals no enrichment of sexually antagonistic polymorphisms on the human X chromosome. Proc. R. Soc. B 289. (DOI:10.1098/rspb.2021.2314).

[35] Harper, J. A. & Morrow, E. H. 2025 The adaptive value of recombination in resolving intralocus sexual conflict by gene duplication. Proc. R. Soc. B 292. (DOI:10.1098/rspb.2024.2629).

[36] Harper, J. A., Janicke, T. & Morrow, E. H. 2021 Systematic review reveals multiple sexually antagonistic polymorphisms affecting human disease and complex traits. Evolution 75, 3087–3097. (DOI:10.1111/evo.14394).

[37] Arnold, S. J., Bürger, R., Hohenlohe, P. A., Ajie, B. C. & Jones, A. G. 2008 Understanding the evolution and stability of the G-matrix. Evolution 62, 2451–2461.

[38] Wyman, M., Stinchcombe, J. & Rowe, L. 2013 A multivariate view of the evolution of sexual dimorphism. J. Evol. Biol. 26, 2070–2080.

[39] Gerdts, E. & Regitz-Zagrosek, V. 2019 Sex differences in cardiometabolic disorders. Nat. Med. 25, 1657–1666. (DOI:10.1038/s41591-019-0643-8).

[40] Wiese, C. B., Avetisyan, R. & Reue, K. 2023 The impact of chromosomal sex on cardiometabolic health and disease. Trends in Endocrinology & Metabolism 34, 652–665. (DOI:10.1016/j.tem.2023.07.003).

[41] Zhernakova, D. V., Sinha, T., Andreu-Sánchez, S., Prins, J. R., Kurilshikov, A., Balder, J.-W., Sanna, S., Franke, L., Kuivenhoven, J. A., Zhernakova, A., et al. 2022 Age-dependent sex differences in cardiometabolic risk factors. Nature Cardiovascular Research 1, 844–854. (DOI:10.1038/s44161-022-00131-8).

[42] Mauvais-Jarvis, F. 2024 Sex differences in energy metabolism: natural selection, mechanisms and consequences. Nature Reviews Nephrology 20, 56–69. (DOI:10.1038/s41581-023-00781-2).

[43] Gibbs, R. A., Belmont, J. W., Hardenbol, P., Willis, T. D., Yu, F. L., Yang, H., Ch’ang, L.-Y., Huang, W., Liu, B. & Shen, Y. 2003 The international HapMap project.

[44] McVean, G. A. & Altshuler, D. M. & Durbin, R. M. & Abecasis, G. R. & Bentley, D. R. & Chakravarti, A. & Clark, A. G. & Donnelly, P. & Eichler, E. E. & Flicek, P., et al. 2012 An integrated map of genetic variation from 1,092 human genomes. Nature 491, 56–65. (DOI:10.1038/nature11632).

[45] Bulik-Sullivan, B., Finucane, H. K., Anttila, V., Gusev, A., Day, F. R., Loh, P.-R., Duncan, L., Perry, J. R. B., Patterson, N., Robinson, E. B., et al. 2015 An atlas of genetic correlations across human diseases and traits. Nat. Genet. 47, 1236–1241. (DOI:10.1038/ng.3406).

[46] Bulik-Sullivan, B. K., Loh, P.-R., Finucane, H. K., Ripke, S., Yang, J., Patterson, N., Daly, M. J., Price, A. L. & Neale, B. M. 2015 LD Score regression distinguishes confounding from polygenicity in genome-wide association studies. Nat. Genet. 47, 291–295.

[47] Hansen, T. F. & Houle, D. 2008 Measuring and comparing evolvability and constraint in multivariate characters. J. Evol. Biol. 21, 1201–1219. (DOI:10.1111/j.1420-9101.2008.01573.x).

[48] Cheverud, J. 1996 Quantitative genetic analysis of cranial morphology in the cotton-top (Saguinus oedipus) and saddle-back (S. fuscicollis) tamarins. J. Evol. Biol. 9, 5–42.

[49] Cheverud, J. M. & Marroig, G. 2007 Comparing covariance matrices: Random skewers method compared to the common principal components model. Genet. Mol. Biol. 30, 461–469.

[50] Team, R. C. 2026 R: A Language and Environment for Statistical Computing. (

[51] Gower, J. C. & Zeilman, B. 1998 Orthogonality and its approximation in the analysis of asymmetry. Linear Algebra and its Applications 278, 183–193. (DOI:10.1016/S0024-3795(97)10090-8).

[52] Zielman, B. 2022 asymmetry: Multidimensional Scaling of Asymmetric Proximities. (

[53] Gosden, T. P., Shastri, K.-L., Innocenti, P. & Chenoweth, S. F. 2012 The B-matrix harbors significant and sex-specific constraints on the evolution of multicharacter sexual dimorphism. Evolution 66, 2106–2116. (DOI:10.1111/j.1558-5646.2012.01579.x).

[54] Watanabe, K., Taskesen, E., van Bochoven, A. & Posthuma, D. 2017 Functional mapping and annotation of genetic associations with FUMA. Nature Communications 8, 1826. (DOI:10.1038/s41467-017-01261-5).

[55] Chen, E. Y., Tan, C. M., Kou, Y., Duan, Q., Wang, Z., Meirelles, G. V., Clark, N. R. & Ma’ayan, A. 2013 Enrichr: interactive and collaborative HTML5 gene list enrichment analysis tool. BMC Bioinformatics 14, 128. (DOI:10.1186/1471-2105-14-128).

[56] Kuleshov, M. V., Jones, M. R., Rouillard, A. D., Fernandez, N. F., Duan, Q., Wang, Z., Koplev, S., Jenkins, S. L., Jagodnik, K. M., Lachmann, A., et al. 2016 Enrichr: a comprehensive gene set enrichment analysis web server 2016 update. Nucleic Acids Res. 44, W90–W97. (DOI:10.1093/nar/gkw377).

[57] Merla, G., Brunetti-Pierri, N., Micale, L. & Fusco, C. 2010 Copy number variants at Williams-Beuren syndrome 7q11.23 region. Hum Genet 128, 3–26. (DOI:10.1007/s00439-010-0827-2).

[58] Lefebvre, P. & Staels, B. 2021 Hepatic sexual dimorphism — implications for non-alcoholic fatty liver disease. Nature Reviews Endocrinology 17, 662–670. (DOI:10.1038/s41574-021-00538-6).

[59] Goossens, G. H., Jocken, J. W. E. & Blaak, E. E. 2021 Sexual dimorphism in cardiometabolic health: the role of adipose tissue, muscle and liver. Nature Reviews Endocrinology 17, 47–66. (DOI:10.1038/s41574-020-00431-8).

[60] Mittendorfer, B. 2005 Sexual Dimorphism in Human Lipid Metabolism1. The Journal of Nutrition 135, 681–686. (DOI:10.1093/jn/135.4.681).

[61] Jayakumar, S. & Hasan, G. 2018 Neuronal Calcium Signaling in Metabolic Regulation and Adaptation to Nutrient Stress. Frontiers in Neural Circuits Volume 12 - 2018. (DOI:10.3389/fncir.2018.00025).

[62] Toprak, U. Lipid Metabolism in Relation to Calcium Homeostasis. (pp. 1–21. Cham, Springer International Publishing.

[63] Asunción-Alvarez, D., Palacios, J., Ybañez-Julca, R. O., Rodriguez-Silva, C. N., Nwokocha, C., Cifuentes, F. & Greensmith, D. J. 2024 Calcium signaling in endothelial and vascular smooth muscle cells: sex differences and the influence of estrogens and androgens. American Journal of Physiology-Heart and Circulatory Physiology 326, H950–H970. (DOI:10.1152/ajpheart.00600.2023).

[64] Koek, W., Campos-Obando, N., van der Eerden, B., De Rijke, Y., Ikram, M., Uitterlinden, A., Van Leeuwen, J. & Zillikens, M. 2021 Age-dependent sex differences in calcium and phosphate homeostasis. Endocrine connections 10, 273–282.

[65] Harris, A., Siggers, P., Corrochano, S., Warr, N., Sagar, D., Grimes, D. T., Suzuki, M., Burdine, R. D., Cong, F., Koo, B.-K., et al. 2018 ZNRF3 functions in mammalian sex determination by inhibiting canonical WNT signaling. PNAS 115, 5474–5479. (DOI:doi:10.1073/pnas.1801223115).

[66] Deshpande, G., Nouri, A. & Schedl, P. 2015 Wnt Signaling in Sexual Dimorphism. Genetics 202, 661–673. (DOI:10.1534/genetics.115.177857).

[67] Hernandez Gifford, J. A. 2015 The role of WNT signaling in adult ovarian folliculogenesis. Reproduction 150, R137–R148. (DOI:10.1530/rep-14-0685).

[68] Xue, R., Lin, W., Sun, J., Watanabe, M., Xu, A., Araki, M., Nasu, Y., Tang, Z. & Huang, P. 2021 The role of Wnt signaling in male reproductive physiology and pathology. Mol. Human Reprod. 27. (DOI:10.1093/molehr/gaaa085).

[69] Murillo-Garzón, V. & Kypta, R. 2017 WNT signalling in prostate cancer. Nature Reviews Urology 14, 683–696. (DOI:10.1038/nrurol.2017.144).

[70] Lombardi, A. P. G., Royer, C., Pisolato, R., Cavalcanti, F. N., Lucas, T. F. G., Lazari, M. F. M. & Porto, C. S. 2013 Physiopathological aspects of the Wnt/β-catenin signaling pathway in the male reproductive system. Spermatogenesis 3, e23181. (DOI:10.4161/spmg.23181).

[71] Yeh, M. M., Shi, X., Yang, J., Li, M.Fung, K.-M. & Daoud, S. S. 2022 Perturbation of Wnt/β-catenin signaling and sexual dimorphism in non-alcoholic fatty liver disease. Hepatology Research 52, 433–448. (DOI:10.1111/hepr.13754).

[72] Ellegren, H. & Parsch, J. 2007 The evolution of sex-biased genes and sex-biased gene expression. Nature Reviews Genetics 8, 689–698. (DOI:10.1038/nrg2167).

[73] Parsch, J. & Ellegren, H. 2013 The evolutionary causes and consequences of sex-biased gene expression. Nature Reviews Genetics 14, 83–87. (DOI:10.1038/nrg3376).

[74] Grath, S. & Parsch, J. 2016 Sex-Biased Gene Expression. Annu. Rev. Genet. 50, 29–44. (DOI:10.1146/annurev-genet-120215-035429).

[75] Khramtsova, E. A., Davis, L. K. & Stranger, B. E. 2019 The role of sex in the genomics of human complex traits. Nature Reviews Genetics 20, 173–190.

[76] Kassam, I., Wu, Y., Yang, J., Visscher, P. M. & McRae, A. F. 2019 Tissue-specific sex differences in human gene expression. Hum. Mol. Genet. 28, 2976–2986. (DOI:10.1093/hmg/ddz090).

[77] Lopes-Ramos, C. M., Chen, C.-Y., Kuijjer, M. L., Paulson, J. N., Sonawane, A. R., Fagny, M., Platig, J., Glass, K., Quackenbush, J. & DeMeo, D. L. 2020 Sex Differences in Gene Expression and Regulatory Networks across 29 Human Tissues. Cell Reports 31, 107795. (DOI:10.1016/j.celrep.2020.107795).

[78] Karlebach, G., Veiga, D. F. T., Mays, A. D., Chatzipantsiou, C., Barja, P. P., Chatzou, M., Kesarwani, A. K., Danis, D., Kararigas, G., Zhang, X. A., et al. 2020 The impact of biological sex on alternative splicing. bioRxiv, 490904. (DOI:10.1101/490904).

[79] Hartman, R. J. G., Huisman, S. E. & den Ruijter, H. M. 2018 Sex differences in cardiovascular epigenetics—a systematic review. Biology of Sex Differences 9, 19. (DOI:10.1186/s13293-018-0180-z).

[80] Wijchers, P. J. & Festenstein, R. J. 2011 Epigenetic regulation of autosomal gene expression by sex chromosomes. Trends Genet. 27, 132–140. (DOI:10.1016/j.tig.2011.01.004).

