## Supplementary Text and Figures for "In search of intralocus sexual conflict in the multivariate and local genetic architecture of metabolic traits in humans"

#### Supplementary File

### Supplementary Note

#### 1) Genome-wide (global) genetic covariances/correlations

We estimated all the SNP-heritabilities, genetic covariances and correlations using LDSC regression method [1, 2]. LD-score of SNP *j* is the sum of the LD correlation of the *jth* SNP with all the other SNPs ($l_j = \sum_k r_{jk}^2$) , where *k* is the total number of SNPs. The method is based on the fact that GWAS effect size of a given SNP includes the effect of all other SNPs it is in LD with [1, 2]. SNPs in high LD will have higher $\chi^2$-statistic on average than SNPs with low LD [1]. The regression of the mean $\chi^2$-statistic on the LD-scores gives SNP-heritability as the slope. SNP-heritability ($h^2_{SNP}$) is the proportion of phenotypic variance explained by the SNPs included in the reference panel (in our case SNPs with MAF between 1% to 50%). As phenotypic variance ($V_P$) is standardized to 1 in LDSC, the $h^2_{SNP}$ is equivalent to the additive genetic variance ($V_A$) [1].

LDSC uses Z-scores from GWAS summary statistics to estimate genetic correlation between traits and sexes. Let *S* denote a set of *M* SNPs, $y_1$ and $y_2$ the phenotypes, *β* and *γ* are the vectors of SNP effect size. Then heritability explained by SNPs in *S* is $h_S^2(y_1) := \sum_j \beta_j^2$ and the genetic covariance among SNPs in *S* is $\rho_S(y_1, y_2) := \sum_{j \in S} \beta_j \gamma_j$.

Under a polygenic model, the expected value of the product of Z-scores, $z_1$ and $z_2$, from two studies for the same SNP *j* is –

$$E[z_1 z_2 l_j] = \frac{\sqrt{N_1 N_2 \rho_g}}{M} l_j + \frac{\rho N_S}{\sqrt{N_1 N_2}} \qquad (1)$$

Where $N_i$ is the sample size of study *i*, $\rho_g$ is the genetic covariance, $l_j$ is the LD score, *M* is the number of SNPs in the reference panel, $N_s$ is the number of overlapping individuals in both the studies, and *ρ* is the phenotypic correlation among number of $N_s$ overlapping samples. The genetic covariance is estimated by regressing $z_{1j}z_{2j}$ on $\frac{l_j}{\sqrt{N_{1j}N_{2j}}}$ and then multiplying the slope by *M*, the number of SNPs in the reference panel (with MAF between 1% to 50%). The intercept ($\frac{\rho N_S}{\sqrt{N_1 N_2}}$), on the other hand, includes the effect of bias and confounding due to sample overlap and population stratification [2].

Genetic correlation between *trait 1* and *trait 2* (or males and females ) was calculated as $r_g = \frac{\rho_g}{\sqrt{h_1^2 h_2^2}}$, where $\rho_g$ is the genetic covariance between traits 1 and 2 (males and females) , and $h_1^2$ and $h_2^2$ are the heritabilities of the two traits (two sexes).

#### 2) Local genetic correlations

We estimated the local genetic correlations using ρ-HESS [3].

The estimator for local genetic covariance at the *ith* region is:

$$\rho_{g,local,i} = Cov\left[\boldsymbol{x}_i^T \boldsymbol{\beta}_i, \boldsymbol{x}_i^T \boldsymbol{\gamma}_i\right] = \boldsymbol{\beta}_i^T \boldsymbol{V}_i \boldsymbol{\gamma}_i \qquad (5)$$

Where ***x*** is the vector of SNPs, ***β*** is the vector of GWAS effect sizes of trait 1, ***γ*** is the vector of GWAS effect sizes of trait 2, and **V** is the LD matrix in the *ith* region. Shared samples across GWASs introduce bias and inflation. Cross-sex-cross-trait genetic correlations are not affected by sample overlap bias because male and female GWAS were conducted in non-overlapping individuals (no individual can be both male and female). The total genome was partitioned into 1703 independent LD-blocks (loci) of approximately 16.4 Mb long, among which 10 loci in chromosome 6 starting from 25684587 to 37572596 bp represent the extended MHC region. One locus in chromosome 2 from 89154526 to 95326452 bp included no SNPs in most of the traits. These 11 regions were removed from the analyses of the local regions. Genetic correlations were obtained by standardizing local genetic covariance estimates by the square root of the corresponding local SNP heritabilities. A total of 1691 loci were analysed, and multiple testing correction was applied using a Bonferroni adjusted threshold of $P < 2.96 \times 10^{-5}$ (0.05/1,691).

### Supplementary Figures

##### Figure 1

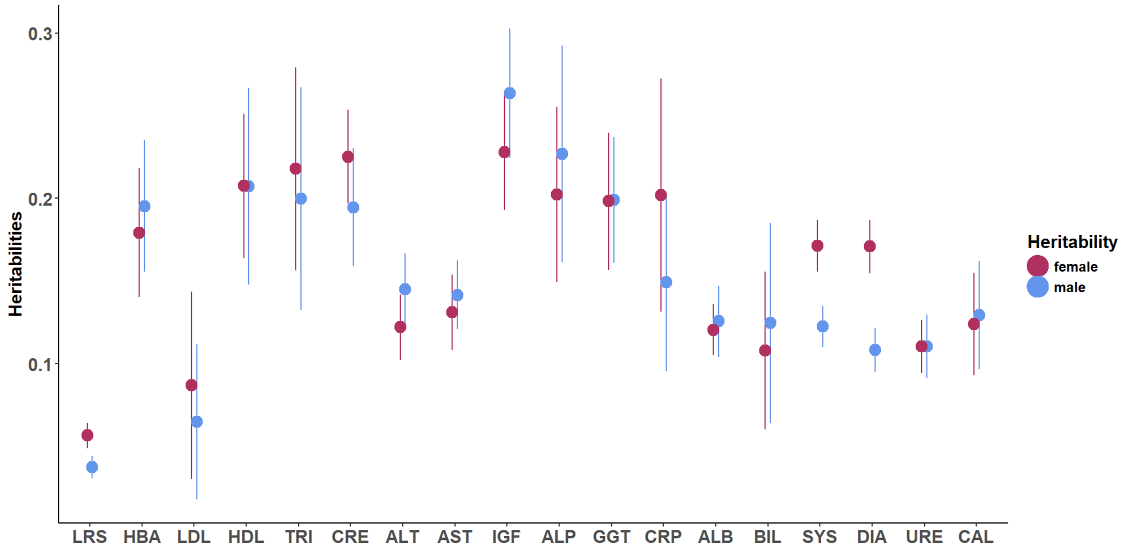

**Figure 1: Sex-stratified SNP-heritabilities of 18 traits.** Male (maroon) and female (blue) SNP-heritabilities along with 95% confidence intervals are plotted for all the metabolic and physiological traits including LRS. Most of the heritabilities are not significantly different between the sexes with LRS, systolic (SYS) and diastolic (DIA) blood pressures showing sex differences.

##### Figure 2

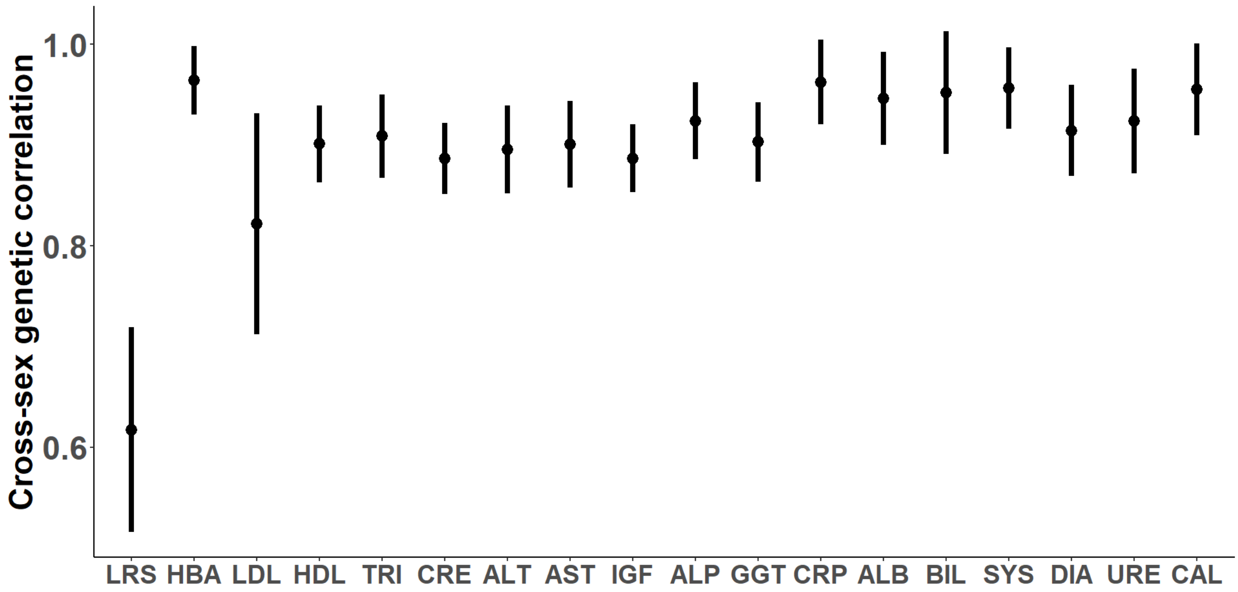

**Figure 2: Cross-sex genetic correlation of 18 traits.** Genetic correlation between male and female traits ($r_{mf}$) with 95% confidence intervals. LRS has the lowest correlation while the other traits having a correlation ~ > 0.8.

#### Figure 3

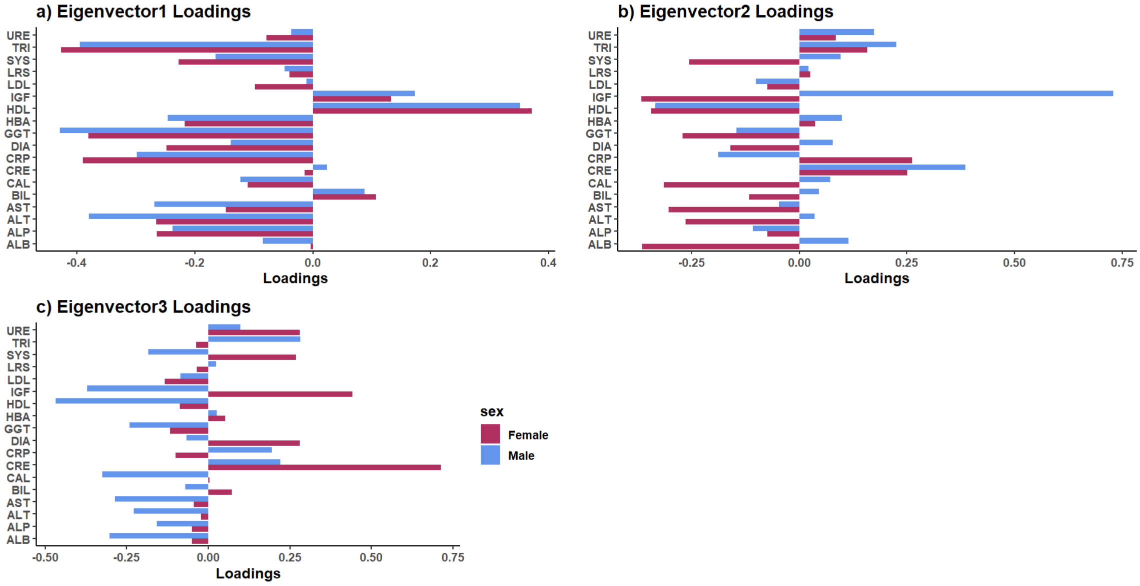

**Figure 3: Loadings of the first three eigenvectors of $G_m$ and $G_f$.** The trait loadings of the first three eigenvectors of male and female G matrices are plotted. a) Loadings of the first eigenvector ($\mathbf{g}_{max}$) are almost similar for the traits in both the sexes, with high loadings for triglycerides (TRI), HDL and GGT. Higher loadings are noted for CRP in females and ALT in males. b) The second eigenvector had asymmetric loadings in the two sexes with IGF1 being highest in males and albumin (ALB), HDL, and calcium (CAL) high in females. c) The third eigenvector also showed difference in trait loadings between the sexes with creatinine (CRE) loading heavily in females and HDL in males.

#### Figure 4

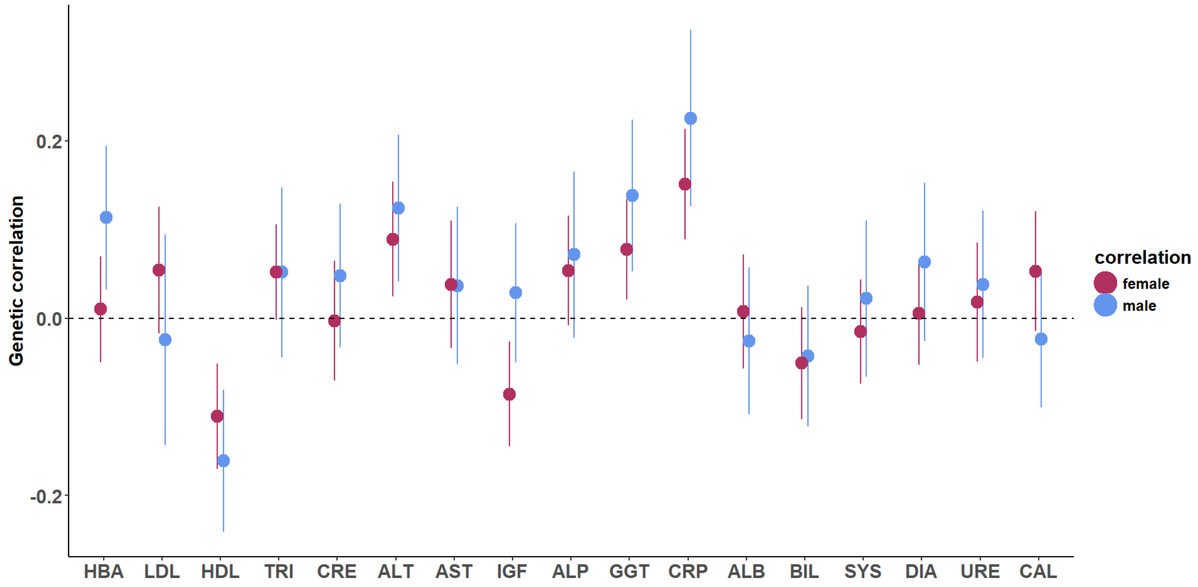

**Figure 4: Trait-fitness genetic correlations.** Genetic correlation between LRS and 17 metabolic and physiological traits in males and females along with 95% confidence intervals. HbA1c (HBA) was significantly correlated only in males and IGF1 only in females. The other traits showing significant correlation in both the sexes were HDL, ALT, GGT and CRP. The rest of the traits did not show any significant trait-fitness correlation

#### Figure 5

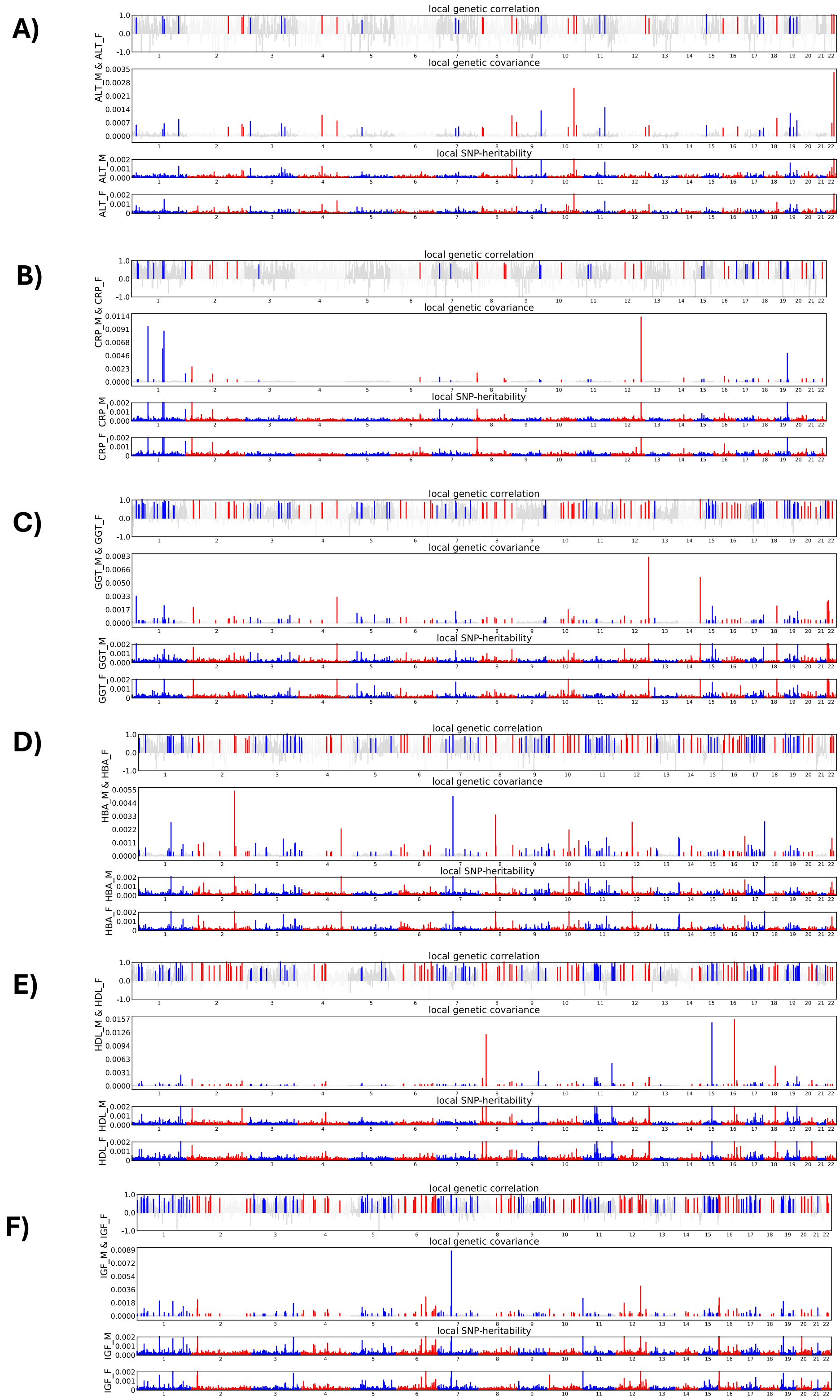

**Figure 5: Local cross-sex genetic covariances and correlations.** Cross-sex correlations between A) male and female ALT, B) male and female CRP, C) male and female GGT, D) male and female HBA, E) male and female HDL, and F) male and female IGF. X-axis contains the chromosomes from 1-22, and Y-axis contains the local genetic covariance/correlations. Only significant regions after Bonferonni correction are represented in colour with red for even chromosomes and blue for odd ones. Lower panels contain the local SNP-heritabilities of the trait-pairs.

**Figure 6**

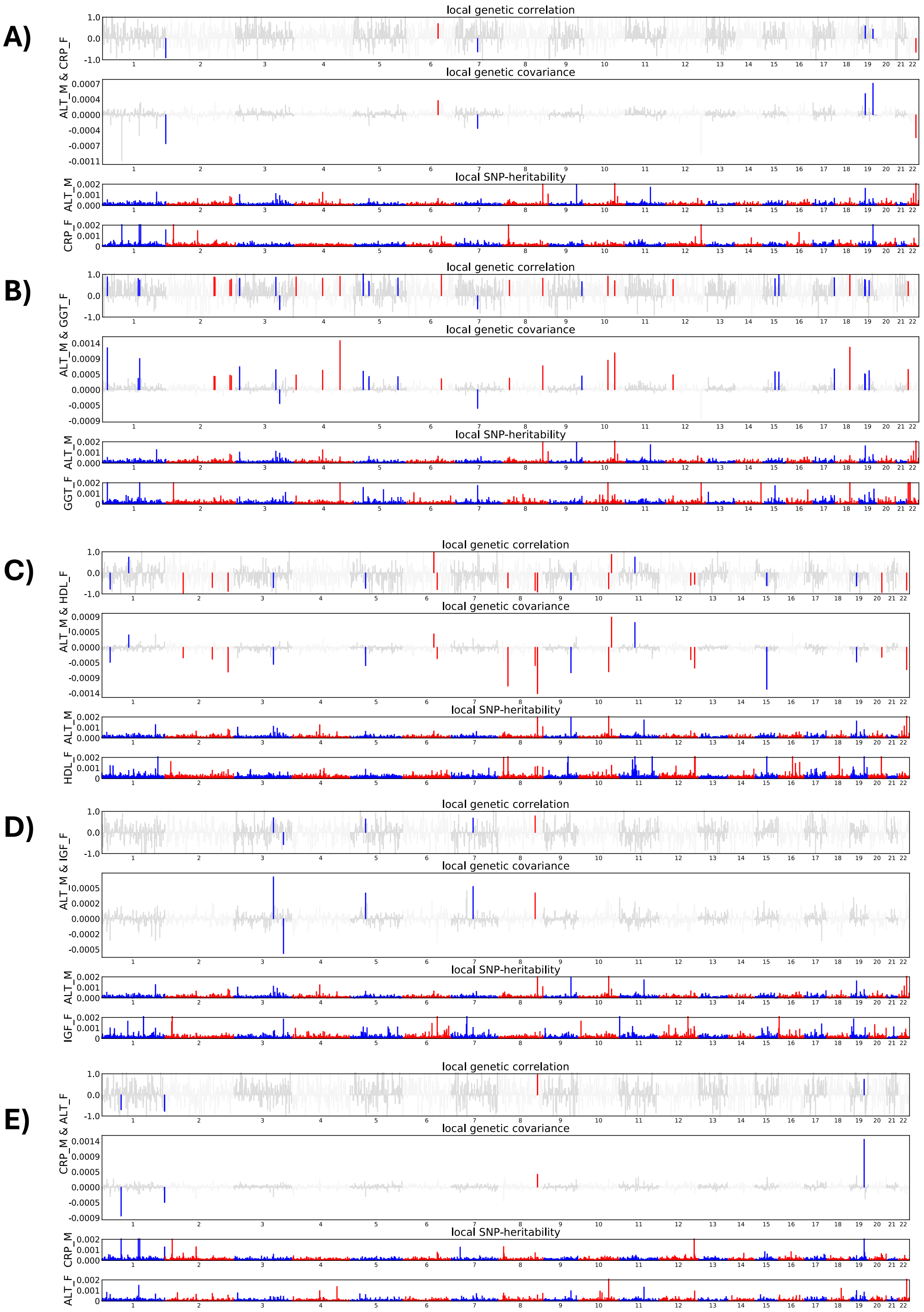

**Figure 6: Local cross-sex-cross-trait genetic covariances and correlations.** Cross-sex-cross-trait genetic genetic correlations between male ALT and A) female CRP, B) female GGT, C) female HDL, and D) female IGF, and E) male CRP and female ALT. Significant regions are marked in colour, red for even chromosomes and blue for odd chromosomes. The local regions showed both positive and negative correlations despite the global genetic correlations being positive or negative.

**Figure 7**

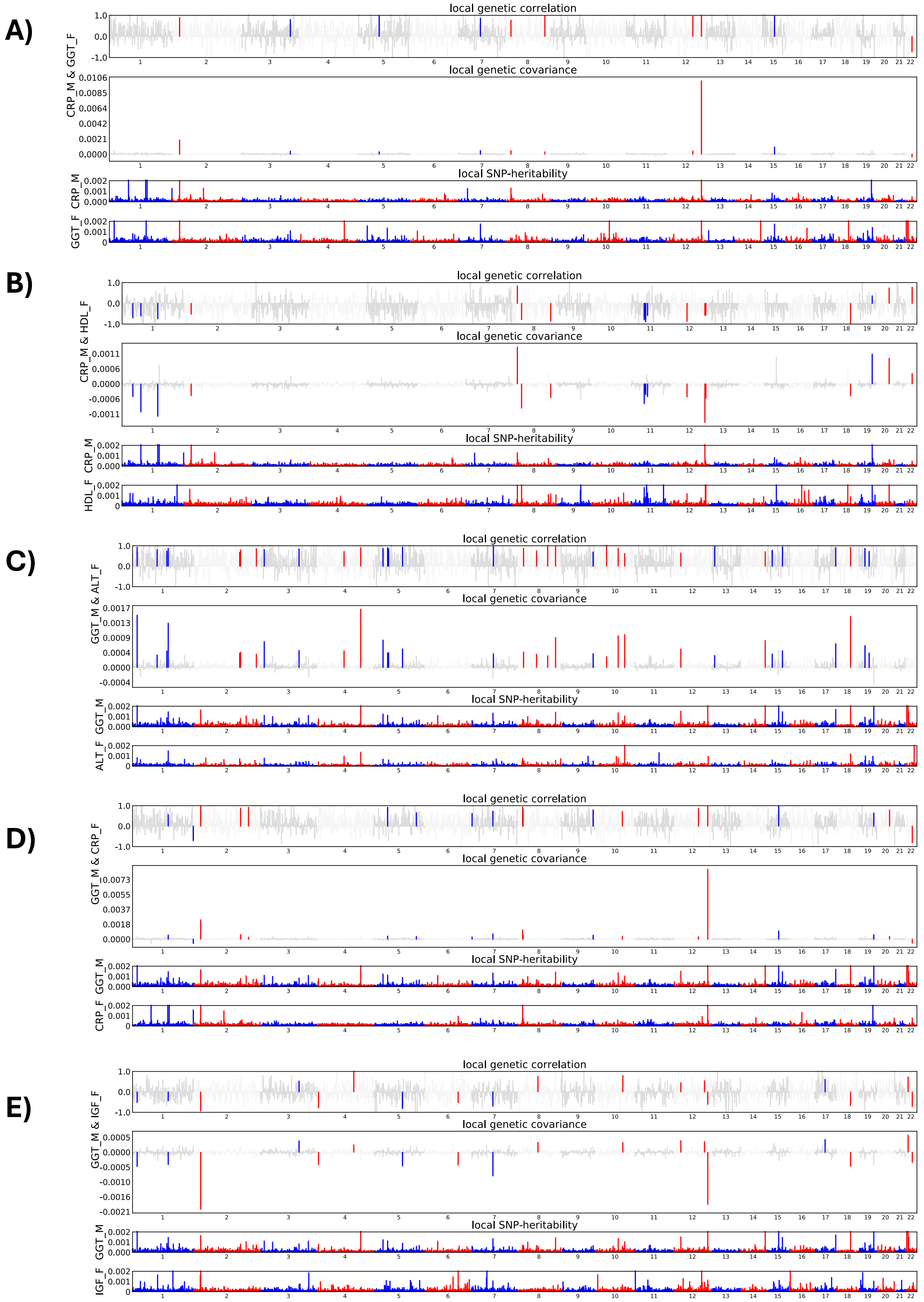

**Figure 7: Local cross-sex-cross-trait genetic covariances and correlations.** Correlations between A) male CRP and female GGT, B) male CRP and female HDL, C) male GGT and female ALT, D) male GGT and female CRP, E) male GGT and female IGF. Significantly correlated regions are highlighted in colour, red for even chromosomes and blue for odd ones.

**Figure 8**

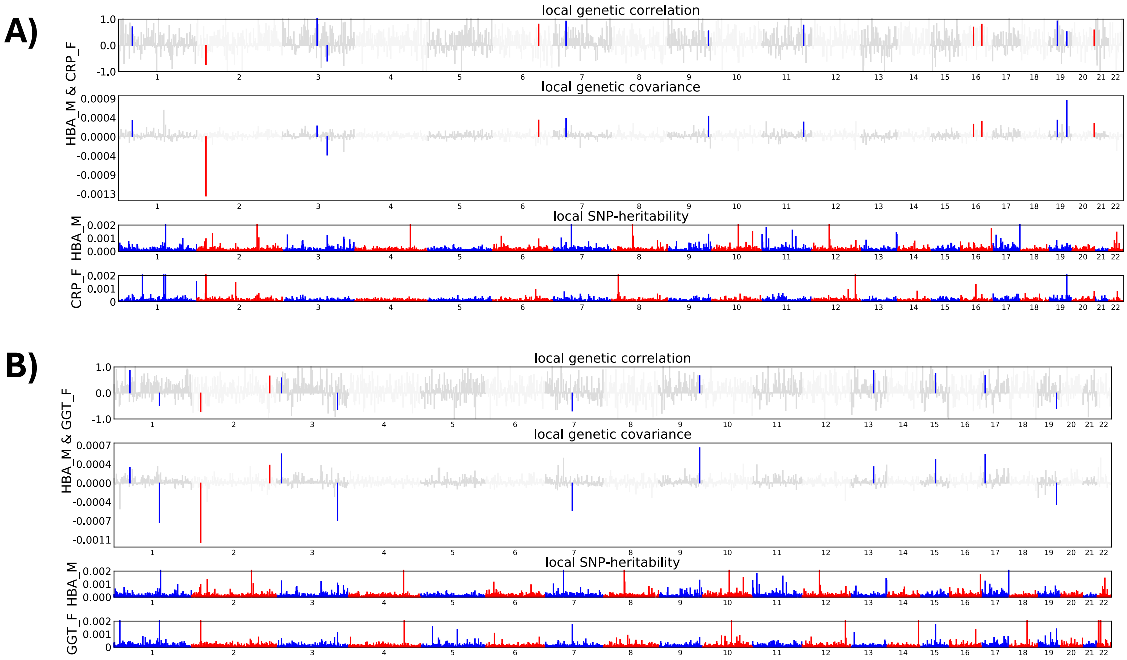

**Figure 8: Local cross-sex-cross-trait genetic covariances and correlations.** Local correlations between A) male HBA and female CRP, and B) male HBA and female GGT. Significantly correlated regions are represented in colour, red for even and blue for odd chromosomes.

**Figure 9**

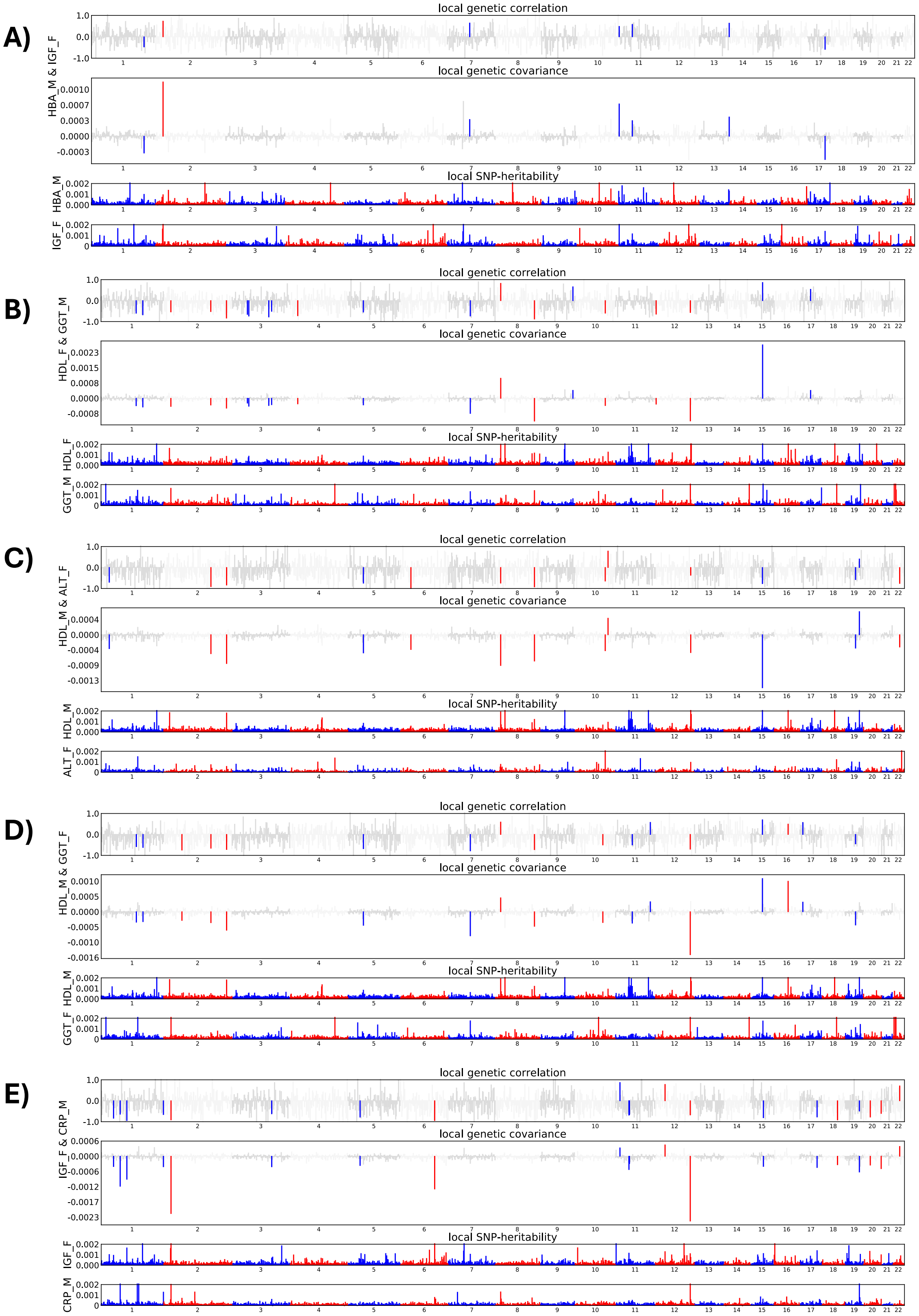

**Figure 9: Local cross-sexcross-trait genetic covariances and correlations.** Local correlations between A) male HBA and female IGF, B) male GGT and female HDL, C) male HDL and female ALT, D) male HDL and female GGT, E) male CRP and female IGF. Significantly correlated regions are represented in colour, red for even chromosomes and blue for odd.

### Representative codes for estimating genome-wide and local correlations

We provide representative scripts for LDSC and ρ-HESS analyses here. Identical pipelines were applied across all trait pairs, across sexes differing only in input summary statistics.

**Code for munging GWAS summary statistics with LDSC**:

munge_sumstats.py --sumstats ~/path/to/file/filename --chunksize 500000 --out output filename --merge-alleles HM3 SNPlist --a1 effect allele column --a2 other allele column --snp rsid --p name of p value column --N sample size --info-min 0.9 --maf-min 0.01 --frq minor_AF --info info

Where chunksize is the number of the SNPs processed by LDSC at a time, merge alleles is the flag for the file name with HapMap3 SNPS which were only used for the analyses, a1 is the flag for the effect allele of GWAS column, a2 is the flag for other allele of GWAS, snp is the flag for the SNP column, p is for pvalue, N for sample size, info-min is the flag of the minimum INFO score of the SNP, maf-min is the minimum minor allele frequency used as a cut off, and frq is the flag for the minor allele column.

**Code for estimating genome-wide genetic correlation with LDSC:**

ldsc.py --rg alt_m.sumstats.gz,ggt_f.sumstats.gz --ref-ld-chr eur_w_ld_chr/ --w-ld-chr eur_w_ld_chr/ --out alt_m_ggt_f

where –rg is the flag for genetic correlation between two munged summary statistics files, --ref-ld-chr is the flag for LD-score files used as independent variable in the regression, --w-ld-chr is the flag for LD-score files to be used for regression weights, --out is the flag for output file.

**Code for estimating local genetic covariance with ρ-HESS**

1) for chrom in {1..22} ;do python hess.py --local-rhog munge_hba_m_chr_bp_nomissing.tsv munge_tri_f_chr_bp_nomissing.tsv --chrom $chrom --bfile ~/hess_noexmhc/Plink_files/1kg_eur_1pct_chr${chrom} --partition ~/hess_noexmhc/EUR/fourier_ls-chr${chrom}.bed --out output_hba_m_tri_f_noexmhc/hba_m_tri_f_step1;done

   Where –local-rhog is the flag for the summary statistics between which correlation is to be estimated, --chrom is the chromosome number, --bfile is the 1000 Genomes European reference files in PLINK format, --partition is the flag for partitioned regions based on LD blocks, --out is the output file.

2) python hess.py --prefix hba_m_tri_f_step1_trait1 --reinflate-lambda-gc 1.0 --out hba_m_tri_f_step2_trait1

   python hess.py --prefix hba_m_tri_f_step1_trait2 --reinflate-lambda-gc 1.0 --out hba_m_tri_f_step2_trait2

   These are the steps to estimate local SNP-heritability for the two traits adjusting for lambda gc which is a correction used for population stratification

3) python hess.py --prefix hba_m_tri_f_step1 --local-hsqg-est hba_m_tri_f_step2_trait1.txt hba_m_tri_f_step2_trait2.txt --reinflate-lambda-gc 1.0 1.0 --num-shared 0 --pheno-cor 0 --out hba_m_tri_f_step3

Step to estimate the local genetic covariance. As these are cross-sex-cross-trait genetic covariances, the number of shared samples (--num-shared) is 0. Consequently, the phenotypic correlation parameter was set to 0, as no sampling covariance due to sample overlap is expected.

#### References

[1] Bulik-Sullivan, B. K., Loh, P.-R., Finucane, H. K., Ripke, S., Yang, J., Patterson, N., Daly, M. J., Price, A. L. & Neale, B. M. 2015 LD Score regression distinguishes confounding from polygenicity in genome-wide association studies. *Nat. Genet.* **47**, 291-295.

[2] Bulik-Sullivan, B., Finucane, H. K., Anttila, V., Gusev, A., Day, F. R., Loh, P.-R., Duncan, L., Perry, J. R. B., Patterson, N., Robinson, E. B., et al. 2015 An atlas of genetic correlations across human diseases and traits. *Nat. Genet.* **47**, 1236-1241. (DOI:10.1038/ng.3406).

[3] Shi, H., Mancuso, N., Spendlove, S. & Pasaniuc, B. 2017 Local Genetic Correlation Gives Insights into the Shared Genetic Architecture of Complex Traits. *The American Journal of Human Genetics* **101**, 737-751. (DOI:10.1016/j.ajhg.2017.09.022).
